# Svirlpool: Multi-sample structural variant calling for Oxford Nanopore sequencing data using local read consensus assembly

**DOI:** 10.1101/2025.11.03.686231

**Authors:** Vinzenz May, Till Hartmann, Dieter Beule, Manuel Holtgrewe

**Affiliations:** Translational Bioinformatics, Institute of Health at Charité Berlin, Luisenstrasse 65, 10115, Berlin, Germany; Medizinische Genetik Mainz, Limbach Genetics eGbR, Haifa-Allee 38, 55128, Mainz, Germany

**Keywords:** structural variants, long-read sequencing, Oxford Nanopore, local assembly, multi-sample calling, Mendelian consistency

## Abstract

**Motivation:** Long-Read Sequencing (LRS), and Oxford Nanopore Technologies (ONT) in particular, has greatly improved the detection of structural genome variants (SVs). Fast alignment-based ONT callers achieve strong benchmark performance, but they necessarily reduce the read sequence to alignment-derived signals when deciding whether variants are shared across samples. This can be limiting for cohort and clinical analyses, especially for insertions and repeat regions where sequence representation matters. We present *Svirlpool*, a multi-sample SV caller for ONT data that builds local consensus assemblies of candidate SV regions and retains the assembled sequence up to the final joint-calling step, where merging tolerances are scaled by a reference-independent noise estimate derived from the reads.

**Results:** We validated Svirlpool on two ONT family datasets: the recent high-quality HG002 Ashkenazi trio and the older Platinum Pedigree family, using the Genome in a Bottle and T2TQ100 benchmarks on the GRCh38, GRCh37, and CHM13v2 references and the Mendelian consistency of native multi-sample calls. We compare against current native joint callers and post-hoc merging workflows. Svirlpool produces highly Mendelian-consistent insertion calls in trio analyses (95.2% on GRCh38 and 95.1% on CHM13v2 at 30x), and on CHM13v2 it reaches the highest insertion and deletion consistency among all tested approaches. Sawfish and Sniffles achieve the highest SV benchmark F1 scores on recent high-quality ONT data, whereas Svirlpool enters the competition with more conservative SV calls. Svirlpool features native, sequence-aware joint calling with retained local consensus sequences and shows a very high Mendelian consistency with sequencing data from different batches and chemistries, which is a common situation in clinical application. Availability and Implementation: Source code, container images, and documentation available at https://github.com/bihealth/svirlpool.

## 1. Introduction

Genomic structural variants (SVs) are causative for a wide array of rare diseases (Collins et al., 2020), Alzheimer’s disease (Vialle et al., 2025), and play an important role in cancer biology (ICGC/TCGA Pan-Cancer Analysis of Whole Genomes Consortium, 2020; Cortés-Ciriano et al., 2020; Wang et al., 2020). SVs are commonly defined as DNA sequence changes of at least 50 base pairs length. Due to the repetitive and segmentally duplicated architecture of the human genome, SV detection remains challenging (Liu et al., 2024). Longer reads can span difficult regions in the genome and carry more structural context, whereas short-read approaches still miss a substantial fraction of insertions and deletions (Chaisson et al., 2019), although more recent studies report improvements (Choo and Imieliński, 2023).

Long-read SV callers can be broadly grouped into: (1) alignment-based methods that extract SV signals from read-to-reference alignments; (2) de novo assembly approaches that first assemble contigs and then compare them to a reference; and (3) local-assembly methods that assemble subsets of reads anchored by initial mappings. De novo assembly can achieve high accuracy but typically requires very high coverage (50x to 90x), and is computationally expensive (Ahsan et al., 2023; Logsdon et al., 2020). Alignment-based methods are fast and perform best with high quality reference genomes but alignment ambiguities in challenging regions can introduce biases that propagate to the results, particularly for Oxford Nanopore Technologies (ONT) reads. The local-assembly strategy offers a middle ground by assembling reads in targeted regions to reduce coverage requirements. A few such methods have been published and validated with Pacific Biosciences data, but, to the best of our knowledge, none for Nanopore data (Denti et al., 2022; Kronenberg et al., 2024).

Long reads from Pacific Biosciences (PB) and Oxford Nanopore (ONT) show different biases in their error profiles, especially given the PacBio HiFi circularized high-accuracy reads (Mikheenko et al., 2022; Park et al., 2025; Sacristán-Horcajada et al., 2021). Key factors for such sequencing errors are tandem repeats, homopolymer runs, and low sequence complexity regions.

In rare disease and cancer studies, SVs are frequently compared across individuals to identify shared variants. However, in repetitive or homologous regions, read alignments may not localize SVs unambiguously, necessitating heuristics to determine variant identity across samples. Existing strategies either offer a multi-sample mode with a built-in merging of variants across samples (Smolka et al., 2024) or refine called variants and then attempt to merge SVs (Kirsche et al., 2023). Recent benchmarks can be found in (Aydin et al., 2025; Helal et al., 2024).

To our knowledge, there are currently three SV callers that have explicit multi-sample SV calling modes: Sniffles2 (PB and ONT) is purely alignment-based and discards read sequence detail for speed, while Sawfish and SVDSS generate local assemblies, but focus on PacBio and therefore ignore the ONT specific biases (Denti et al., 2022; Saunders et al., 2025). Moreover, post hoc merging tools such as Survivor and Jasmine (Jeffares et al., 2017; Kirsche et al., 2023) provide practical ways to combine single-sample calls and can be highly effective when the underlying calls are of high quality. They do not, however, retain local read consensus sequence or use a read-derived local noise model when deciding whether nearby calls represent the same variant.

We developed Svirlpool, a local-assembly SV caller that is tuned for ONT data with an explicit multi-sample joint-calling mode to find shared variants across individuals from different sequencing runs. Svirlpool generates local consensus assemblies to capture sequence context, then performs SV comparison and merging using tolerance parameters informed by local sequence features, including sequence complexity, k-mer sketch similarity, and overlap with annotated tandem repeats, as well as noise patterns generated from the original read sequences. To reduce ONT-specific artifacts, Svirlpool filters SV calls driven by short homopolymer alterations and short GT-rich or AT-rich subsequences. This design aims to provide robust single- and multi-sample SV analysis in challenging regions with per-sample genotype annotations, while accepting the higher computational cost of local consensus assembly.

## 2. Methods

### 2.1. Method Description

#### 2.1.1. Overview

Figure 1 gives an overview of the algorithm implemented in Svirlpool. The input is the alignments of one or more samples together with the reference sequence and some annotation tracks (see Section 2.1.6). The output is a VCF file (The 1000 Genomes Project Consortium, 2019) of structural variants with their genotypes and technical/quality annotations in each sample.

**Figure 1.**
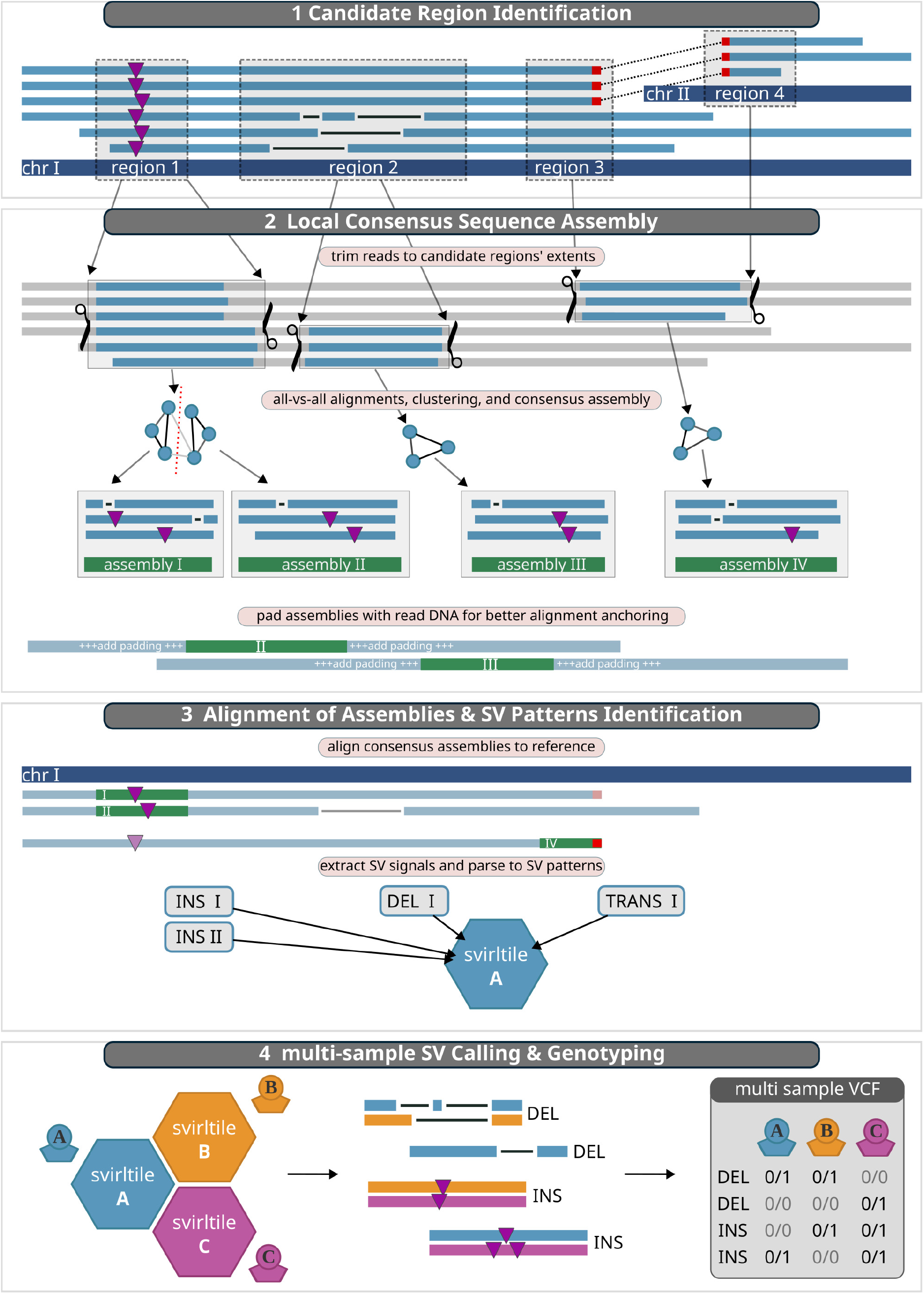
Svirlpool Workflow: 1. Signals from read alignments are collected to construct candidate regions. 2. Reads are trimmed according to the bounds of the candidate regions, respecting connected regions via break ends of underlying reads. The trimmed sequences are aligned all versus all and based on a distance measure grouped with spectral clustering. Each cluster of trimmed reads is assembled and padded with flanking raw read sequences. 3. All padded consensus assemblies are aligned to the reference genome. SV signals are extracted from the consensus alignments and parsed to SV pattern objects, which are stored in a database file, called “svirltile”. 4. All svirltiles are combined in a multi-sample SV processing step, where the SV patterns are merged into SV composites, which contain the signals from different consensus sequences and samples. SV calls are generated from the SV composites and genotyped and written to a multi-sample VCF file.

As is common with complex algorithms, Svirlpool consists of a hierarchy of steps. The highest level consists of four *parts* described in the following sections. Parts 1–3 run independently for each individual sample whereas part 4 runs on multiple samples at once.

Svirlpool scores pairs of signals with a similarity score and normalizes the scores by the local depth of coverage. The algorithm then takes strong signals as seeds for candidate regions and attempts to merge signals into clusters for candidate regions supporting a putative SV based on the similarity scores. The merging takes into consideration the distance on the reference, the signal sizes, and the signal’s overlap with annotated tandem repeats. Svirlpool stores each candidate region together with the supporting signal, read names, and read alignment intervals. When two or more candidate regions are supported by the same read via break-ends (BNDs), we call them connected, and the downstream algorithm parts process them together.

#### 2.1.2. Candidate region identification (Part 1)

The algorithm scans over the read alignments of a single sample. The alignments produced by the read mapper contain insertions/deletions (*indels*) within the alignments (*intra-alignment signal*). Further, two local alignments of adjacent sequence in the same read may show interspersed alignments or align on opposite strands in the reference (*inter-alignment signal*). Inter-alignment signal implies break-ends of the sample’s genome with respect to the reference.

Svirlpool collects intra-alignment signals (indels) using empirically determined thresholds. It then filters the signal using provided annotation tracks and heuristics for alignment and reference artifacts observed empirically in ONT data. The algorithm computes depth of coverage tracks from filtered data, for normalization purposes, and it computes a copy number track that is later used to guide the read clustering process to approximate the number of clusters by the expected zygosity. The copy number and depth tracks are again used in the final genotyping step.

We assume that true SVs are expressed in the read alignments by similar, proximal signals. Therefore, the task for SV identification becomes separating the true SV signals from noise, i.e. introduced by sequencing errors, incorrect reference sequence, or alignment artifacts.

Each signal that passes a two-threshold filter (minimum absolute strength and minimum coverage-normalized strength) seeds a flanking interval whose width depends on signal type: a fixed radius (default 300 bp) for break-end signals, and 10% of the signal size (minimum 50 bp) for insertion and deletion signals, so that larger events naturally capture more context. Annotated tandem repeat regions are appended as additional seeds to guarantee coverage of known repetitive loci, and all intervals are sorted and merged with a configurable buffer radius using bedtools merge to form *proto-candidate regions*. Proto-CRs whose read count exceeds 6*×* the genome-wide median are discarded as likely artefactual pile-ups (e.g. in segmental duplications). Adjacent surviving proto-CRs are further merged when their gap is within twice the smaller of their respective median indel sizes (capped at 300 bp) and their insertion and deletion signal size distributions are sufficiently similar (Cohen’s *d <* 0.5); this secondary merge captures split seeds that share a common SV origin. Finally, each region is extended symmetrically to a minimum size (default: 1,200 bp) to provide sufficient flanking context for the downstream read-trimming and assembly steps.

#### 2.1.3. Local read consensus assembly (Part 2)

Svirlpool considers each connected group of candidate regions separately. The algorithm first **collects all read alignments** of each group. It then **trims** the reads to the candidate regions’ extents, generating a **pool *P* of *cut reads*** which is used as the input for the subsequent **clustering step**.

In the idealized case, a candidate region group stems from a single SV in (without loss of generality) heterozygous state and, except for sequencing errors, all reads align perfectly to one of the two distinct alleles. However, real data shows different challenges including: errors in the reference, ambiguities in the sequence alignments, copy number variable regions, high heterogeneity of structural variants ranging from VNTRs over simple to complex structural variants. Thus, we designed our clustering algorithm to address these challenges with minimal assumptions on SV types and focusing on sequences instead. The clustering first attempts a fast separation of reads by their summed insertion and deletion signals inside the candidate regions. If this does not yield compact and well-separated clusters, Svirlpool falls back to all-vs-all alignments of the trimmed reads (each pair of reads is aligned independently, not jointly as in a multiple sequence alignment). These alignments are parsed into insertion, deletion, and break-end signal profiles. A pairwise distance is computed from the size-damped signal sizes, optional positional importance-density weights, read-length differences, and interior break-end evidence, and is transformed into a similarity matrix with a Gaussian kernel. The reads are then grouped with spectral clustering on this precomputed similarity matrix after removing length and connectivity outliers. The number of clusters is bounded by the local copy number estimation, which runs the Viterbi algorithm on fixed size bins of median coverages with fixed transition probabilities of 90%, no state change and an equal distribution of state change probabilities. The bin size is scaled with the median read alignment length. Mathematical details and parameter values are provided in the Supplementary Methods.

Next, Svirlpool computes a **consensus assembly from each read cluster**. For this, it uses the lamassemble software (Frith et al., 2021). All trimmed reads of one cluster are then aligned to their consensus assembly to determine the interval each read covers on the consensus, and to save any occurring insertion or deletion signals to estimate the local noise distribution in the structural variant calling step.

For each consensus assembly, Svirlpool stores the *core consensus* sequence from the cut read assemblies as well as flanking *padding sequence*. The much shorter consensus core sequence is not necessarily unambiguously mappable due to its reduced length. Our solution is to add flanking DNA letter sequence from the one or two reads that extend the farthest from the core interval. This padding stabilizes the genome-wide alignment of the consensus assembly sequence and preserves flanking context but is ignored for later variant extraction.

#### 2.1.4. From SV primitives to SV patterns (Part 3)

In this part, Svirlpool first aligns the padded consensus sequences to the target reference using minimap2. It then considers the alignments of the core consensus sequences and parses insertions, deletions, and break ends of this core interval into **SV primitives**.

The parsing works as follows: contiguous insertions/deletions are interpreted as insertion (INS)/deletion (DEL) SVs. Large clipped or discordant split segments are interpreted as unresolved novel adjacencies (a.k.a. break-ends/BNDs). Svirlpool stores these SV primitives together with technical annotations such as identifier of sample, consensus sequence ID, and depth of coverage. It further annotates them with overlapping tandem repeat annotations. Also, Svirlpool performs an estimation of the local position noise/distortion by counting the insertion and deletion signals from the trimmed reads to their consensus, weighted by their exponential distance.

The rationale behind the noise estimation is our observation that while some loci show highly concordant SV signals, others show a more distorted signal. Such regions are characterized by low sequence complexity, which leads to Nanopore sequencing errors and general alignment ambiguities, often expressed in many small insertion and deletion signals instead of compact single events. Most such regions are found in and around VNTRs. Svirlpool does not fully characterize VNTRs, but instead reports them as insertions or deletions, and allows the subsequent analysis of the local consensus assemblies for more detailed characterizations of low complexity regions.

The next step is **converting the SV primitives into SV patterns**. It converts primitives from the same consensus assembly alignment into higher-level SV patterns using pre-defined topologies. Svirlpool uses a simple pattern matching algorithm without probabilistic interpretation to avoid overinterpretation. The resulting higher-level SV patterns represent SVs like insertions, deletions, inversions, and complex re-arrangements. SV patterns contain their original SV primitives and offer an interface to query properties of the determined SV types, e.g. length of the inserted sequence.

A common special case is a mixed set of insertions and deletions which are frequent in alignments within tandem repeats. These sets are reduced to a single direction by the net signed length (insertions/deletions correspond to positive/negative net lengths). Such interspersed insertions and deletions inside annotated tandem repeats are merged within the consensus sequence of the same sample (*horizontal merging*) to a single insertion or deletion SV pattern.

The Svirlpool algorithm converts intra-alignment insertion/deletion SV patterns into simple insertions and deletions. It parses break ends to identify large insertions and deletions from inter-alignment signals. Translocations and inversions are identified from break ends as well. In the case of solitary break ends lacking adjacency, they are emitted as VCF unpaired break ends. Finally, any residual unused break ends are left to be complex patterns. This preserves the raw structure for refinement by external downstream tools rather than forcing a classification that is most likely wrong.

#### 2.1.5. Structural variant calling on multiple samples (Part 4)

This part consolidates the annotated per-sample SVs into final, genotyped cohort calls. This part scales from a single sample well to ten samples, but possibly more.

At first a per-sample **horizontal merging** handles the case that two SV patterns from the same consensus sequence express the same SV pattern, but one is constructed from intra-alignment signals (INS/DEL) and the other from break ends. This horizontal merging step is different from the horizontal merging of the SV patterns, where only INS and DEL signals within repeats were merged. As described above, this commonly occurs in the case of tandem repeats or other challenging reference regions. The merging takes one or more SV patterns and merges them into one SV *composite* putatively describing the same underlying variant. Each SV composite is a container of its underlying SV patterns. Most SV composites, however, are created from a single SV pattern.

After performing merging within the signals of each single consensus sequence of one sample, Svirlpool performs the **vertical merging of SV composites** across original consensus sequences and samples into multi-sample *composites*. For this, it compares the distance of the signals on the reference genome, the size of the SVs, and a k-mer based similarity measure. The distance and size comparisons are scaled by the noise distortions from the aligned trimmed reads, and by the sequence complexity of the inserted or deleted sequences, if feasible.

Genotypes are determined per sample at each structural variant locus using a binomial likelihood model. Variant-supporting (alt) and total reads overlapping the locus are identified from precomputed coverage interval trees. For deletions, reads are queried in 100 bp windows around both breakpoints rather than across the full span to avoid missing supporting reads. The local copy number (CN) is obtained from copy-number tracks when available, defaulting to CN=2 (diploid). For each candidate genotype the expected alternate-allele fraction is the number of alternate copies divided by the copy number, clamped to the interval [*ϵ*, 1 − *ϵ*] with a fixed error rate *ϵ* = 0.05 that absorbs read noise and reference over-counting. For a diploid locus the expected fractions are therefore 0.05 (0/0), 0.5 (0/1), and 0.95 (1/1); for CN=1, 0.05 (ref) and 0.95 (alt); and for CN=3, 0.05, 1*/*3, 2*/*3, and 0.95. Genotype likelihoods are computed as the binomial probability of the observed alternate-read count given the total coverage and the expected fraction, then normalized to posterior probabilities assuming uniform priors. The most likely genotype is selected, and a Phred-scaled genotype quality (GQ), capped at 60, is reported alongside total coverage, reference read count, and variant read count. As a final, sequence-aware correction, when a single candidate region yields two or more distinct consensus assemblies for the same sample, the corresponding locus is necessarily heterozygous, and any homozygous-alternate call for that sample is overridden to heterozygous.

Records are flagged `PASS` if at least one sample has three or more supporting reads; otherwise, they are flagged `LowQual`. Breakpoint precision is approximated by repeat content: `PRECISE` if none of the contributing primitives overlap annotated repeats, otherwise `IMPRECISE`.

#### 2.1.6. Genome annotation data

In addition to the samples’ alignments in BAM format (Li et al., 2009), Svirlpool needs the reference in FASTA format with annotations in BED format (Kent et al., 2002). We use the annotations by Pacific Biosciences, public with the PBSV software for **tandem repeats** (see External Resources section). Together with our software, we provide a script to generate all genome regions of **mononucleotide runs** of length 6 and more (Pockrandt et al., 2020; Smit et al., 2013). Extended information on the annotation data we used for this manuscript can be found in the External Resources section.

#### 2.1.7. Implementation and practical notes

Svirlpool’s source code is made available as open source under the MIT license and can be found on GitHub together with instructions on how to run it on a sample dataset: https://github.com/bihealth/svirlpool. The software was implemented in Python 3.12 using the following libraries: biopython v.1.83, intervaltree v.3.1.0, matplotlib v.3.8.4, numpy v.1.26.4, pandas v.2.2.2, plotly v.5.22.0, pysam v.0.22.1, scipy v.1.13.1. Further external software used: lamassemble v.1.7.2, samtools v.1.20, mosdepth 0.3.8, bedtools v.2.32.1, bcftools v.1.20. Between different parts of the algorithm, data is persisted as lightweight SQLite database files allowing fast downstream lookup. Svirlpool expects ONT alignments in BAM or CRAM format produced by the minimap2 aligner (version 2.2 and above) and creates VCF v. 4.2 files (The 1000 Genomes Project Consortium, 2019). A detailed explanation of the output can be found on the GitHub page referred to above. We used Svirlpool version 0.2.0 to generate all presented results.

### 2.2. Benchmark and evaluation

#### 2.2.1. Data

We used two publicly available real-world Nanopore sequencing datasets. First, the Ashkenazi Jewish trio, with the child HG002, the father HG003, and the mother HG004. The trio was sequenced with the ligation-sequencing-kit-v14 using the SQK-LSK114 protocol and base called with Dorado v0.8.3 with model dna_r10.4.1_e8.2_400bps@v5.0.0. The sequencing reads data are publicly and freely available in uBAM format (see External Resources section). To test the batch effect strength, we added two more variants of the HG002 sample: a 2022 R9.4.1 dataset basecalled with Guppy v. 6.3.7, and a 2023 R10.4.1 dataset basecalled with Dorado v. 0.3.0. Full sequencing specifications, basecalling parameters, alignment commands, and subsampling procedures for these two additional datasets are given in the Supplementary Methods.

We aligned the already basecalled ultra-long ONT reads to HG38, HS1, and HG19 with minimap2 (version 2.30-r1287) and arguments: `-x map-ont`. Second, the Platinum Pedigree family, of which 10 samples from three generations in six trios are publicly available, were all sequenced with an older setup: flowcell FLO-PRO002 R9.4.1 and sequencing kit SQK-ULK001, then basecalled with Guppy v. 6.3.7. The exact sequencing specifications can be found in Kronenberg et al. (2024). We aligned the reads to HG38 and HS1 with minimap2 (version 2.28-r1209) and arguments: `-x map-ont`.

We subsampled all sequencing data to 30x, 20x, 10x, and 5x to approximate situations closer to our expectations in the clinical and research application based on the average depth of coverage that was calculated with mosdepth excluding secondary alignments by setting the minimum mapq to 1 (version 0.3.8: mosdepth `--no-per-base --fast-mode --mapq 1`) and samtools (version 1.20: `samtools view --subsample fraction --subsample-seed depth`), with ‘fraction’ chosen to approximate the target coverage and as seed we chose the target coverage to have different subsets of reads after each subsampling.

#### 2.2.2 SV-calling evaluation

Sniffles (v. 2.7.2) offers inbuilt multi-sample variant calling that facilitates weak signals across samples to increase the sensitivity in such cases. We know of no other method that is validated on Nanopore data with this feature, so we compared our results with Sniffles results. Furthermore, we compared to the other local-assembly method Sawfish (Saunders et al., 2025), although it is not tuned for ONT data, but Pacific Biosciences HiFi data, which features lower per base error rates and different error profiles. We also attempted to include FocalSV, but could not generate completed whole-genome or reduced benchmark-region callsets despite a dedicated workflow with FocalSV-compatible alignments and high-memory runs; details are given in the Supplementary Methods.

We benchmarked Sniffles, Sawfish, and Svirlpool with the Genome in a Bottle (GiaB) v5.0q SV benchmark for HG002 (for GRCh38, GRCh37, CHM13v2). We additionally ran the T2TQ100-v1.1 SV benchmark, and for GRCh37, excluding Sawfish, the v0.6 (legacy) benchmark (Zook et al., 2020). We compared the truth sets with truvari version 5.4.0 (English et al., 2022). All tests with truvari were executed with the `--passonly` flag set and the `--sizemin` parameter set to 50, to exclude SVs smaller than 50 bp. All SVs overlapping tandem repeats (TR) in the T2TQ100-v1.1 and V5 benchmarks are neither merged nor left-aligned. This causes many representation-sensitive misses in the benchmarking, so that precision, and especially recall are reduced for all tools in repeat-overlapping regions. We therefore report both full benchmark results and complementary with-TR and without-TR subsets to separate general SV-calling performance from this repeat representation issue. We limited the testing on the legacy v0.6 SV benchmark to the tier 1 (T1) high confidence regions and the canonical chromosome regions (chr1– 22, X, Y). We limited the GRCh38 (HG38) and CHM13v2 (HS1) benchmarks to all canonical chromosome regions and the benchmark regions definitions that are distributed together with the benchmark vcf files.

#### 2.2.3. Mendelian consistency evaluation

The Mendelian consistency tests were applied to both the HG002 trio and the six trios of the Platinum Pedigree family, which consist of ten related individuals. In each trio we force-called SVs, so that genotypes like ‘./.’ were treated as ‘0/0’. The Mendelian consistency test is a module of Svirlpool. It checks if the genotypes can be explained by inheritance or if they violate the possible inheritance patterns. The consistency test is generalized to hemizygous variants, duplications, and triplications.

Sniffles, Sawfish, and Svirlpool each have a native joint SV-calling mode. It typically results in a multi-sample VCF file, which was used to test each trio therein. For each coverage subsample stage (30x, 20x, 10x, 5x), and all reference genomes, we ran the Mendelian test if possible. We also stratified the results along SV type (insertion and deletion) and SV size (bins of 50–100, 100–350, 350–1000, 1k–10k, *>*10k).

#### 2.2.4. HG002 batch and chemistry robustness evaluation

To evaluate whether joint-calling performance degrades when samples from different sequencing chemistries or batches are combined, we formed mixed-chemistry HG002 trios. In these trios the child sample was taken from either the 2022 R9.4.1 dataset (Guppy v. 6.3.7) or the 2023 R10.4.1 dataset (Dorado v. 0.3.0), while both parents were always taken from the 2025 HG002 trio (SQK-LSK114, Dorado v. 0.8.3). All three tools were applied in their native joint-calling modes and the resulting calls were additionally post-hoc merged with Jasmine (version 1.1.5) and Survivor (version 1.0.7). Mendelian consistency was evaluated as described in the previous section across both GRCh38 and CHM13v2 reference genomes and subsampled coverage levels of 5x, 10x, 20x, and 30x. Sawfish was included only where it completed; above 5x coverage it was out-of-memory killed at 1 TB of main memory on these datasets.

#### 2.2.5. Merging methods evaluation

We applied Survivor (version 1.0.7) and Jasmine (version 1.1.5) to the individual SV calls of all single samples to combine them into family VCF files, or a single trio in the case of HG002 with parents. The resulting merged VCF files were subsequently tested for Mendelian consistency as described in the previous section.

## 3. Results

We developed Svirlpool, a local consensus assembly-based germline SV caller, that can work with .bam and .cram alignment files without special aligner flags. It emits records for different types of structural variants, including insertions, deletions, inversions, inverted deletions, inverted insertions, and complex re-arrangements, although the benchmark validation in this study focuses on insertions and deletions. We have tested Svirlpool on whole genome human germline Oxford Nanopore data of different versions and on their alignments to the CHM13v2 (HS1), GRCh38 (HG38), and GRCh37 (HG19) genome references. Svirlpool can perform SV calling on a whole genome, or a defined subset of regions.

### 3.1. Mendelian consistency test results

Figure 2 summarizes the Mendelian consistency results. Across both trios, the native joint-calling modes of Sniffles and Svirlpool reach high consistency, and the higher-quality HG002 trio reads yield notably higher values than the older Platinum Pedigree data. Sub-figure A stratifies by depth of coverage. On the HG002 trio, Svirlpool’s native joint calling reaches the highest overall consistency among the native joint-calling modes (94.4% on GRCh38 at 30x), followed closely by Sniffles (93.5%), with Sawfish substantially lower (87.0%) except at 5x, where Sawfish (81.7%) outperforms both Svirlpool (76.3%) and Sniffles (71.6%). Sawfish could not terminate with up to 1 TB of main memory on the Platinum Pedigree data above 5x coverage, so it is restricted to 5x there.

**Figure 2.**
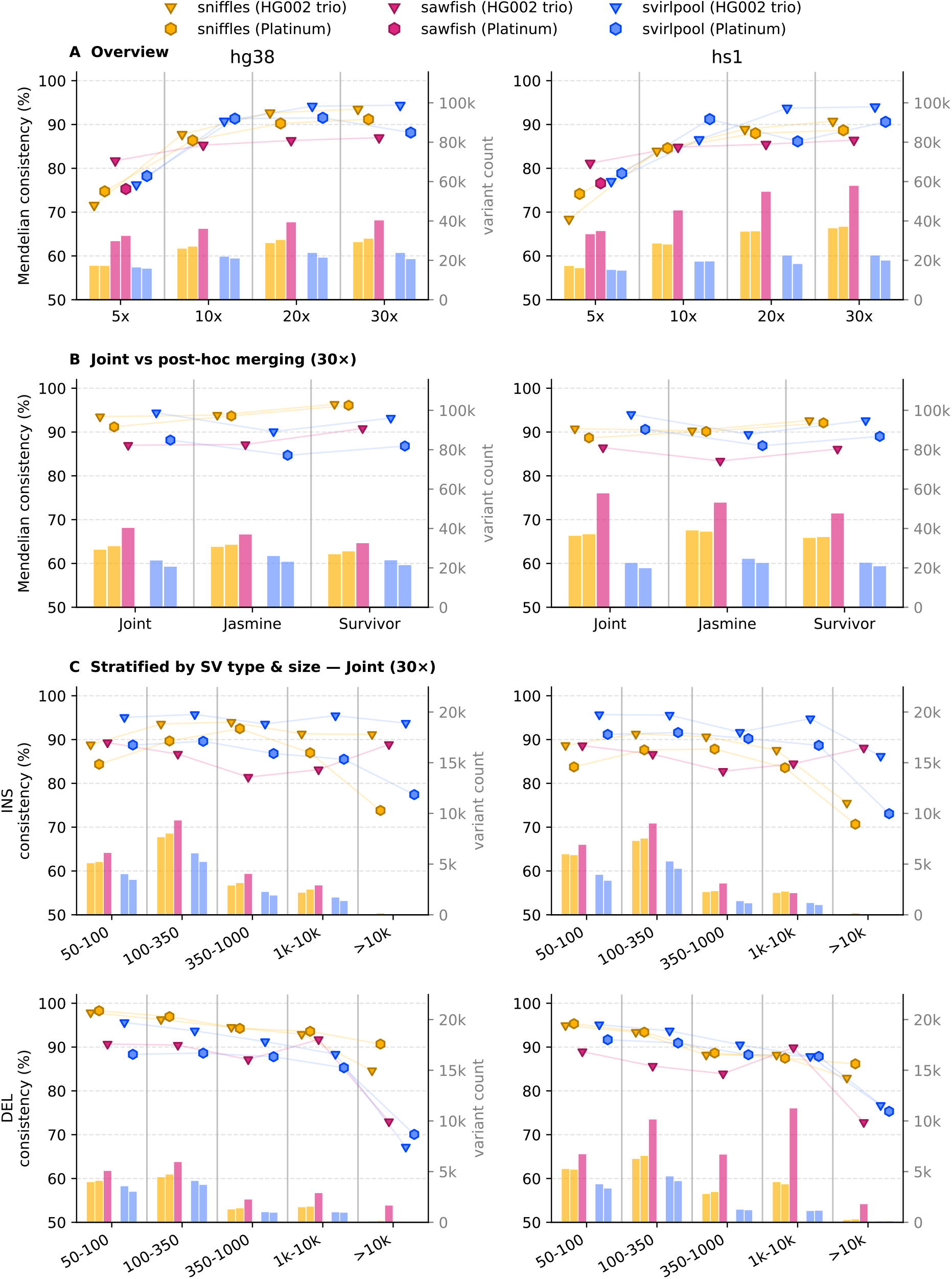
Mendelian consistency results. The three sub-plots show the stratified results of the two datasets: HG002 trio and mean of the six Platinum Pedigree samples. The Mendelian consistency is indicated by the markers and measured by the left vertical axis. The bars show the absolute SV counts, which are measured with the right vertical axis. Note that we were not able to generate Sawfish results for the Platinum samples above 5x depth of alignment coverage due to out-of-memory kills at 1 TB memory. **Sub-figure A** shows the Mendelian consistency stratified across different depth of coverage parameters from 5x to 30x. On the HG002 trio at 30x on GRCh38, Svirlpool reaches 94.4%, Sniffles 93.5%, and Sawfish 87.0%; at 5x, Sawfish (81.7%) outperforms Svirlpool (76.3%) and Sniffles (71.6%). **Sub-figure B** shows the same results for only 30x coverage. They are compared to the 30x results of the post-hoc merging tools Jasmine and Survivor. On GRCh38 the highest overall consistency is achieved by Sniffles merged with Survivor (96.4%), followed by Svirlpool native (94.4%) and Sniffles native (93.5%); Survivor also raises Sawfish from 87.0% to 90.8%. **Sub-figure C** shows the 30x results stratified by SV type (insertions and deletions), and by size. Among native callers, Svirlpool achieves the highest insertion consistency on both references (95.2% on GRCh38, 95.1% on CHM13v2); Sniffles leads on deletions on GRCh38 (96.1%), while Svirlpool leads on both insertion and deletion consistency on CHM13v2 (95.1% and 93.1%, respectively), exceeding all tested approaches including the post-hoc mergers.

Sub-figure B compares the native joint-calling modes against the post-hoc merging tools Jasmine and Survivor on the 30x family data. The single strongest combination for Mendelian consistency is Sniffles single-sample calls merged with Survivor, which reaches 96.6% for insertions and 96.1% for deletions on GRCh38. Survivor markedly improves the Sniffles results, whereas it does not improve, and for insertions slightly degrades, the already-high native results of Svirlpool; the effect on Sawfish is mixed. Importantly, Svirlpool reaches comparable consistency (95.2% for insertions on GRCh38) directly from its native joint-calling step, without any separate post-hoc merging.

Sub-figure C stratifies the 30x results by SV type and size. Among the native joint callers, Svirlpool calls insertions most consistently on both references (95.2% on GRCh38 and 95.1% on CHM13v2), clearly above the native Sniffles values (92.0% and 89.8%, respectively). On the CHM13v2 (T2T) reference Svirlpool attains the highest insertion and deletion consistency of all tested approaches, including the post-hoc mergers. Deletions are most consistent with Sniffles on GRCh38 (96.1% native). Smaller variants are generally called more consistently in the Mendelian sense, while variants larger than 10 kb give mixed results across all tools.

To assess robustness to sequencing chemistry and batch differences, we additionally formed mixed-chemistry HG002 trios in which the child sample was replaced with either the 2022 R9.4.1 or the 2023 R10.4.1 dataset while the parents remained from the 2025 trio. Svirlpool maintains near-identical Mendelian consistency in these cross-chemistry trios: 95.4% for insertions and 94.3% for deletions on GRCh38 at 30x (96.0% and 94.5% on CHM13v2). Sniffles native joint calling is likewise stable at 92.1% insertions and 96.0% deletions on GRCh38 at 30x; post-hoc merging with Survivor raises it to 97.7% and 96.0%, the highest insertion consistency values observed in the batch experiment. Sawfish was out-of-memory killed at 1 TB of main memory on all coverage levels above 5x in these datasets and could therefore not be fully evaluated in the cross-chemistry setting. Full coverage- and size-stratified results, as well as all post-hoc merging comparisons, are shown in Supplementary Figs. S8 and S9.

### 3.2. SV benchmark test results

Figure 3 presents the v5 Genome in a Bottle benchmark limited to the HG002 variants that do not overlap annotated tandem repeats, for the GRCh38 and CHM13v2 reference genomes. Note that the benchmark also contains SV types other than insertions and deletions, so the combined results in the top panels are not simply the sum of the insertion and deletion panels. Sawfish achieves the highest overall F1 score, followed by Sniffles and then Svirlpool: at 30x on GRCh38 the overall F1 scores are 0.95 (Sawfish), 0.95 (Sniffles), and 0.92 (Svirlpool), and on CHM13v2 0.96, 0.96, and 0.93, respectively. The difference is primarily in recall: on GRCh38 at 30x, recall is 0.94 (Sawfish), 0.92 (Sniffles), and 0.89 (Svirlpool), while precision is comparable across all tools (0.95–0.97). The ordering holds when insertions and deletions are considered separately: insertion F1 on GRCh38 at 30x is 0.95 (Sawfish), 0.94 (Sniffles), and 0.90 (Svirlpool); deletion F1 is 0.96, 0.94, and 0.91. On CHM13v2 at 30x, insertion F1 scores are 0.96 (Sawfish), 0.96 (Sniffles), and 0.92 (Svirlpool), and deletion F1 is 0.97, 0.94, and 0.92. Sawfish and Sniffles already reach near-final accuracy at 10x coverage, while 20x and 30x are nearly identical for all tools. Contrary to the Mendelian consistency results, Sniffles performs relatively better on insertions than on deletions in the benchmark.

**Figure 3.**
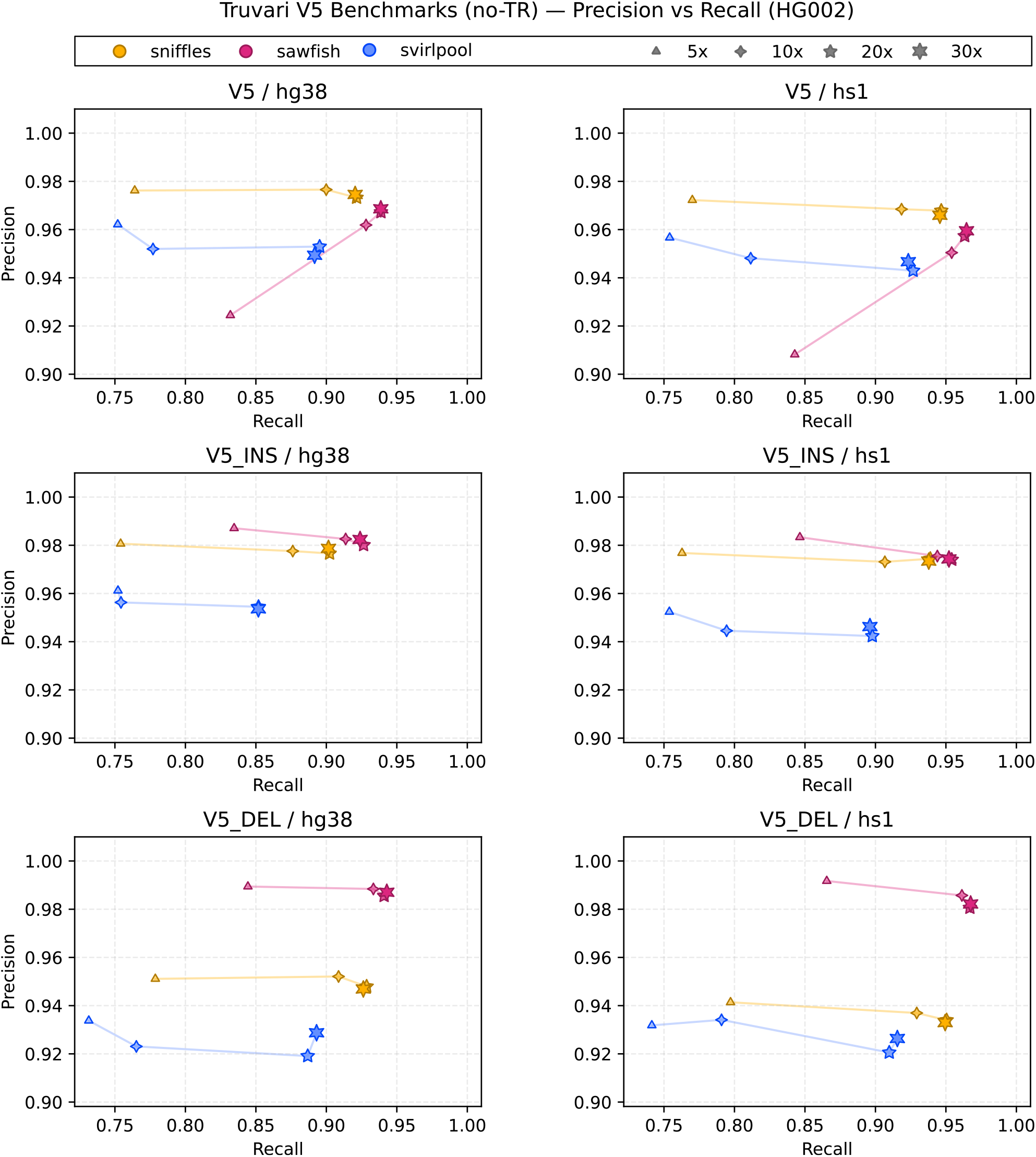
SV benchmark results. This figure shows the results of the v5 Genome in a Bottle SV benchmark limited to SV calls that don’t overlap annotated tandem repeats. The T2TQ100-v1.1 benchmark results are presented separately in Supplementary Fig. S3. Each marker encodes a different depth of coverage from 5x to 30x. We stratified the results along SV type, and reference genome. Note that not only insertions and deletions are part of the benchmark, but other SVs and break ends as well, so that the top plots are not explained by combining the INS and DEL plots. At 30x on GRCh38, overall F1 scores are 0.95 (Sawfish), 0.95 (Sniffles), and 0.92 (Svirlpool); on CHM13v2, 0.96, 0.96, and 0.93. For insertions at 30x on GRCh38, F1 scores are 0.95 (Sawfish), 0.94 (Sniffles), and 0.90 (Svirlpool); for deletions 0.96, 0.94, and 0.91. Sawfish and Sniffles largely plateau after 10x–20x, while Svirlpool shows a larger improvement between 5x and 20x.

When tandem-repeat-overlapping SVs are included, precision and especially recall drop substantially for every tool (Supplementary Fig. S2). As detailed in Section 3.2, this reflects the sensitivity of repeat-region representations: one biological repeat expansion or contraction can be represented either as several nearby indels or as one merged, left-aligned event.

We have further tested and stratified the T2TQ100-v1.1 benchmark, but the results are very similar and follow the pattern of the v5 results, so they are added to Supplementary Fig. S3. The results of the legacy v0.6 benchmark show a similar picture in Supplementary Fig. S4.

The absolute number of calls that pass the quality filter also differs substantially across tools (Supplementary Fig. S5). At 30x coverage Svirlpool produces on average approximately 26 000 PASS calls per sample on GRCh38 and 23 000 on CHM13v2, compared to approximately 36 000 for both Sniffles and Sawfish on GRCh38. This lower call count is consistent with Svirlpool’s conservative assembly-based evidence thresholds and is reflected in its lower recall relative to the other tools.

### 3.3. Read alignment quality

The sequencing samples of the older Platinum Pedigree data and the newer HG002 trio data have significant differences. We measured the alignment gaps per megabase and the alignment fragment lengths. The mean of the median gap counts of the Platinum Pedigree samples is 169.5 and 69.8 in the case of the HG002 trio (Mann-Whitney test: *U* =0, *p*=0.007). The mean of the median fragment lengths of the Platinum Pedigree samples is significantly longer than those of the HG002 trio (Mann-Whitney test: *U* =2, *p*=0.028). More detailed plots can be seen in Supplementary Fig. S1.

### 3.4. Resource consumption

Table 1 summarises the single-sample wall time and peak memory of each tool as the minimum, median, and maximum across all coverage levels and reference genomes for both datasets. Sniffles executes much faster and with much less memory than the local-assembly methods across the whole distribution. Svirlpool is the most time-intensive method and shows the widest memory range, reaching its maximum on the Platinum Pedigree samples aligned to the CHM13v2 (HS1) reference genome, whereas Sawfish reaches its memory maximum on the Platinum Pedigree samples aligned to GRCh38. More detailed visualisations and tables can be found in the supplementary material (Supplementary Figs. S6 and S7, Supplementary Table S1). The resource consumptions of the SV joint-calling steps of all tools (Sniffles, Sawfish, Svirlpool, Jasmine, and Survivor) are generally lower than the single sample processing steps and are visualized and comprehensively expressed in a table in the supplementary material (Supplementary Fig. S6, Supplementary Table S2). Thus, Svirlpool is not intended as the fastest default caller for very large cohorts. Its computational cost is most justified when native sequence-aware joint calling, repeat-aware merging, or access to the local consensus sequences is part of the downstream analysis question.

**Table 1.** Single-sample resource consumption per tool and dataset, given as the minimum, median, and maximum across all completed runs (coverage levels 5x–30x and reference genomes). Wall time in minutes, peak resident memory in gigabytes.

| Tool | Dataset | Wall time (min) |  |  | Peak memory (GB) |  |  |
| --- | --- | --- | --- | --- | --- | --- | --- |
|  |  | Min | Median | Max | Min | Median | Max |
| Sniffles | HG002 trio | 0.9 | 2.7 | 5.2 | 2.4 | 7.9 | 14.1 |
|  | Platinum | 0.9 | 3.4 | 9.7 | 2.7 | 7.6 | 13.3 |
| Sawfish* | HG002 trio | 10.8 | 60.6 | 119.3 | 28.0 | 67.5 | 131.2 |
|  | Platinum | 13.0 | 30.0 | 141.8 | 27.8 | 58.5 | 218.4 |
| Svirlpool | HG002 trio | 140.3 | 305.6 | 856.5 | 9.9 | 46.4 | 279.4 |
|  | Platinum | 67.9 | 554.7 | 1392.2 | 12.7 | 15.7 | 350.5 |

\* Sawfish did not complete on the Platinum Pedigree data above 5x coverage (out-of-memory at 1 TB), so those runs are excluded. See Supplementary Figs. S7 and S8 for full per-run breakdowns and joint-calling resources.

## 4. Discussion

In this study we have investigated how local assembly can contribute to improving SV calling and genotyping in ONT long read data. We introduce the new tool Svirlpool implementing this approach, which we make available as open source under a permissive license. We performed thorough benchmarking and comparison with established best-practice SV calling tools for different linear reference genomes, data sets, and analyzed challenges in different variant classes and benchmark data sets. The established Sniffles best practice tool, especially with the improvements introduced in recent versions >=2.7.2 and working on recent ONT data, remains the most versatile tool, with specific strengths in the tested deletion calling and very low execution times.

Post-hoc merging methods like Jasmine and Survivor show great potential for the identification of shared variants across samples. It is interesting to observe that Sawfish, which was specifically designed for PacBio HiFi data, performs the best on the T2T-derived SV benchmarks, although the Mendelian consistency tests on both the native joint calling and the post-hoc merging results indicate that Sawfish is still not the best choice for multi-sample or cohort analyses of ONT data.

It is worth noting that the SV benchmarking with the GiaB version v5.0q and the T2TQ11-v1.1 SV benchmark structurally favor (local) assembly-based SV callers that don’t merge indels within tandem repeats and don’t left-align them. The convention in newer benchmarks has shifted to represent variants purely as reference-relative deviations. This is however, in our clinical application not always appreciated, because heterogeneous data from different chemistries and batches need to be compared. Therefore, we implemented a feature in Svirlpool to extract the locally assembled sequence for exact sequence-to-reference comparisons, but handle the variant reporting and merging in the native joint calling in the more established way of merging indels within tandem repeats and left-aligning them.

Svirlpool addresses a problem that is central to cohort and clinical studies: identifying the structural variants that several individuals from different batches or even chemistries share, rather than analysing each sample in isolation. It is not the fastest caller and it does not reach the highest benchmark F1 score on recent high-quality ONT data. Its intended role is narrower: native, sequence-aware ONT joint calling with retained local consensus sequences. The established alignment-based approach to multi-sample ONT calling, exemplified by Sniffles, reduces the underlying read sequence to alignment-derived signals for speed when deciding whether two calls represent the same variant. Svirlpool instead retains the assembled sequence throughout the workflow, up to the final joint-calling step. Its methodological core is how the merging decisions are informed: for each local read cluster we record the alignment gaps between the cluster’s reads and their consensus assembly. These gaps quantify a local noise level that derives entirely from the reads and is therefore independent of reference bias, and we use this estimate to scale the distance and size tolerances when merging variants, both within a sample and across samples in the joint-calling step. This is the main reason to use Svirlpool alongside a fast alignment-based caller: the merging decisions use sequence and local read-derived noise rather than reference-derived signals alone. It does not replace variant-database lookups, but it provides a consistent identification of shared variants within a processed cohort.

We assess the joint calling primarily through Mendelian consistency (Section 3.1). On its own, this metric is insufficient: a method that merges aggressively appears highly consistent simply because it collapses distinct variants into shared calls. Mendelian consistency is therefore only informative together with benchmark precision. The single most consistent pipeline we tested is Sniffles single-sample calls merged post-hoc with Survivor, reaching 96.6% for insertions and 96.1% for deletions on GRCh38 while retaining high benchmark precision; this is a genuinely strong and practical combination. Svirlpool’s native joint calling reaches comparable consistency (95.2% for insertions on GRCh38) directly, without a separate merging tool. We cannot quantify possible over-merging from Mendelian consistency alone; instead, the benchmark precision, the lower PASS call counts, and the repeat-stratified analyses provide the main checks that the consistency is not driven only by indiscriminate collapsing of nearby calls.

Insertions remain the most difficult and clinically most important case. As discussed by Kronenberg et al. (2024), detecting and characterizing inserted sequence is hard, and existing methods differ considerably. This is where Svirlpool’s sequence-preserving design has the largest effect: across a trio, its native joint calling resolves insertions more consistently than the native joint calling of Sniffles (95.2% versus 92.0% on GRCh38, and 95.1% versus 89.8% on CHM13v2), and on the CHM13v2 (T2T) reference it attains the highest insertion and deletion consistency of all tested approaches, including the post-hoc mergers.

The cross-chemistry batch experiment (Supplementary Figs. S8 and S9) provides additional evidence for clinical applicability. Svirlpool is highly robust to chemistry and batch mismatches: its consistency in mixed-chemistry trios (95.4% insertions on GRCh38 at 30x) is essentially unchanged from the same-chemistry result (95.2%), confirming that the sequence-aware merging based on local read-derived noise tolerates inter-chemistry variation well. Sniffles combined with Survivor also performs strongly in this setting, reaching 97.7% insertion consistency on GRCh38 at 30x, the highest value in the batch experiment, and should be considered a practical and low-cost alternative when assembly-based joint calling is not required. Sawfish could not be evaluated beyond 5x coverage in the batch datasets due to out-of-memory termination at 1 TB, which limits its use for multi-chemistry cohorts on current hardware.

We examined whether a local-assembly SV caller tuned for PacBio HiFi transfers to ONT, using Sawfish (Saunders et al., 2025). On the most recent, highest-quality ONT data, where gap rates are low and reads are long, Sawfish achieves the best precision and recall (Section 3.2). However, on the older Platinum Pedigree data with higher error rates, Sawfish did not terminate within 1 TB of memory above 5x coverage (Section 3.4), consistent with its design for the very low error rates of HiFi reads. Sawfish is therefore highly informative about how well HiFi-oriented local assembly can transfer to recent ONT data, but we do not treat it as a like-for-like ONT method across all datasets tested here.

The legacy v0.6 GiaB SV benchmark annotated SVs within tandem repeats to the upstream limit of the repeat and merged the insertions and deletions within one tandem repeat and one allele. Svirlpool follows this convention for intra-repeat indel signals by merging them to a single net insertion or deletion and then reporting the event at the repeat-consistent position. The newer benchmark versions (T2TQ100-v1.1 and v5.0q) do not merge and left-align these repeat-internal events. As a result, every tested tool suffers representation-sensitive losses in precision and recall: insertions and deletions within tandem repeats are hard to localize exactly, read aligners tend to produce multiple indel signals in contracted or expanded repeats, and one biological event can appear as many smaller events in the benchmark. The position of a gap can also shift substantially within a repeat given only a few single-nucleotide differences (IGV examples, Supplementary Figs. S10–S13). To show this effect explicitly, we separated both the benchmark and the tool results into subsets with and without tandem-repeat overlap.

We extended the comparison to post-hoc merging by applying Jasmine and Survivor to each caller’s single-sample calls and testing the merged cohorts for Mendelian consistency. Both yielded clear gains for Sniffles, mixed outcomes for Sawfish, and, for Svirlpool, slightly worse results than its native joint calling. Since neither tool performs local assembly, they are methodologically closer to Sniffles than to Sawfish or Svirlpool, and their resource consumption is very low. The strong performance of Sniffles with Survivor shows that, for high-quality single-sample calls, a permissive proximity-based merge can be highly effective; Svirlpool offers an alternative in which comparable, sequence-aware reasoning is applied natively and without reference-anchored tolerances.

Several limitations constrain these conclusions. On high-quality data Svirlpool does not reach the best precision and recall, trailing both Sniffles and Sawfish on the benchmarks, and it is considerably more resource-intensive than alignment-based calling (Section 3.4, Table 1); the read-derived merging signal is also technically demanding to compute. Although Svirlpool emits inversions, inverted insertions and deletions, and complex rearrangements, the benchmarks used here are essentially insertion and deletion truth sets, so accuracy for the other variant types is not established in this work. Our genotyping model (Section 2.1.5) is likewise not separately evaluated for genotype concordance. Finally, Mendelian consistency alone cannot reveal overly tolerant merging; only simulation could, and we did not simulate because the structural phenomena observed in real ONT data could not be reproduced faithfully. Testing, for example, HG002 against unrelated parents from the Platinum Pedigree would not be informative, as no expected truth value is available for comparison.

Finally, the consensus sequences from local assembly hold potential for downstream applications, such as dedicated sequence analysis of VNTRs in genes such as FMR1 (Loomis et al., 2013) and MUC1 (Vrbacká et al., 2025).

## 7. Data availability

All sequencing data used in this study are publicly available (see External Resources section). Source code is available at https://github.com/bihealth/svirlpool.

## 8. Author contributions

V.M. and M.H. conceived the study. V.M. implemented Svirlpool and conducted all experiments. T.H. contributed to benchmarking and evaluation. D.B. and M.H. supervised the project. V.M., T.H., D.B., and M.H. wrote and reviewed the manuscript.

## 9. Acknowledgements

We thank Prof. Dr. Johannes Köster from the University of Duisburg-Essen for his support and valuable discussions.

## 10. External Resources

HG002 trio data: https://epi2me.nanoporetech.com/giab-2025.01/

giab_2025.01/basecalling/hac/HG002/PAW70337/calls.sorted. bam

giab_2025.01/basecalling/hac/HG003/PAY87794/calls.sorted. bam

– giab_2025.01/basecalling/hac/HG004/PAY87778/calls.sorted. bam

HG002 2022 (R9.4.1) reads: https://s3-us-west-2.amazonaws.com/human-pangenomics/index.html?prefix=T2T/scratch/HG002/sequencing/ont/03_08_22_R941_HG002_rebasecalling-guppy-6.3.7/03_08_22_R941_HG002_6.fq.gz

HG002 2023 (R10.4.1) reads: s3://ont-open-data/giab_2023.05/analysis/hg002/hac/PAO89685.pass.cram

Platinum Pedigree sequencing data: https://github.com/Platinum-Pedigree-Consortium/Platinum-Pedigree-Datasets

RepeatMasker tracks — HS1: https://hgdownload.soe.ucsc.edu/goldenPath/hs1/bigZips/;HG38: https://hgdownload.soe.ucsc.edu/goldenPath/hg38/bigZips/;HG19: https://hgdownload.soe.ucsc.edu/goldenPath/hg19/bigZips/

PBSV annotation tracks: https://github.com/PacificBiosciences/pbsv/tree/master/annotations

