## supplementary material for "Svirlpool: Multi-sample structural variant calling for Oxford Nanopore sequencing data using local read consensus assembly"

### S1 Supplementary Methods

#### S1.1 Attempted FocalSV comparison

We attempted to include FocalSV as an additional local-assembly comparison by building a dedicated Snakemake workflow around the public FocalSV scripts. The workflow was restricted to dataset and reference combinations for which the required FocalSV prior resources were available: Genome in a Bottle data on hg19 and hg38, and Platinum Pedigree data on hg38. CHM13v2/hs1 was excluded because no matching FocalSV prior VCF was available. The workflow considered the 5x, 10x, 20x, and 30x subsampled read sets.

For each sample, reads were realigned with minimap2 using FocalSV-compatible ONT settings: `minimap2 --MD -Y -L -a -x map-ont`. Alignments were sorted with samtools and checked for chr-prefixed contig names, because FocalSV expects chromosome identifiers such as `chr1`. Region discovery was run with `0_define_region.py` in ONT mode, using the corresponding reference FASTA and prior VCF. To reduce the downstream workload, the discovered FocalSV regions were intersected with the subsampled HG002 v5 benchmark BED regions before running FocalSV auto mode. The final calling command used `focalsv.py` with `--chr_num 0`, `--target_bed`, `--data_type ONT`, and 64 threads for `--num_cpus`, `--num_threads`, and `--early_threads`. Resource requests were scaled by coverage from a 30x baseline: 64 GB and 2880 min for read alignment, 32 GB and 360 min for region definition, and 320 GB and 4800 min for FocalSV auto calling, with retry-based doubling of these resources.

Despite this reduced target-region workflow and high-memory runs, we did not obtain completed FocalSV callsets suitable for the same whole-genome or benchmark region comparison used for Sniffles, Sawfish, and Svirlpool. We therefore did not include FocalSV in the final quantitative benchmark figures.

#### S1.2 HG002 legacy chemistry datasets

To investigate the effect of sequencing chemistry and batch on joint-calling performance, two additional HG002 datasets produced with older chemistries were included alongside the primary 2025 HG002 trio data.

**2022 R9.4.1 dataset.** Reads were obtained from flowcell 6 of the March 2022 ONT ultra-long sequencing run, using the Oxford Nanopore Ultra-Long DNA Sequencing Kit with R9.4.1 chemistry. Raw reads were basecalled with Guppy v. 6.3.7 using the configuration model `dna_r9.4.1_450bps_modbases_5hmc_5mc_cg_sup_prom.cfg`. Read data are publicly available (see External Resources in the main manuscript). Reads were aligned to GRCh38 with minimap2 and coordinate-sorted with samtools:

```
minimap2 -t 32 -a -x map-ont -Y GRCh38.fa \
```

```
    03_08_22_R941_HG002_6.fq.gz \
    | samtools sort -@32 -O bam -o HG002.2022.30x.bam
```

Subsampling to 20x, 10x, and 5x target coverage was performed with seqkit, using sampling fractions and seeds tied to the target coverage:

```
seqkit sample -j 64 -t dna -p 0.68 -s 20 \
    -o HG002.20x.fastq.gz HG002.30x.fastq.gz
seqkit sample -j 64 -t dna -p 0.34 -s 10 \
    -o HG002.10x.fastq.gz HG002.30x.fastq.gz
seqkit sample -j 64 -t dna -p 0.17 -s 5  \
    -o HG002.5x.fastq.gz  HG002.30x.fastq.gz
```

**2023 R10.4.1 dataset.** Reads were obtained from the ONT open data release (May 2023), sequenced on a PromethION flow cell (FLO-PRO114M) with the SQK-LSK114 Ligation Sequencing Kit and R10.4.1 chemistry. Basecalling was performed with Dorado v. 0.3.0. The CRAM file is publicly available (see External Resources in the main manuscript). Subsampling to 30x, 20x, 10x, and 5x was performed directly from the CRAM with samtools, excluding the chrEBV contig:

```
samtools view -b -@ 4 \
    --subsample <fraction> \
    -T GCA_000001405.15_GRCh38_no_alt_analysis_set.fna \
    -o PAO89685.tmp.bam PAO89685.pass.cram \
    $(samtools view -H PAO89685.pass.cram | grep "^@SQ" \
      | cut -f2 | sed 's/SN://' | grep -v "chrEBV")
```

where `<fraction>` was adjusted per target coverage level (30x, 20x, 10x, and 5x).

#### S1.3 Svirlpool local consensus assembly details

Svirlpool processes candidate regions as connected candidate-region containers. For each container, it first queries the copy-number track over all candidate region intervals in the container. The maximum copy number is used as an upper bound on the number of read clusters, with a minimum bound of two. Containers with estimated copy number greater than four are skipped for consensus generation, because the local haplotype structure is considered too complex for the current implementation.

For every candidate region in a container, Svirlpool retrieves primary read alignments from the input BAM/CRAM and obtains the corresponding full-length read sequences. Supplementary or hard-clipped read sequence is resolved through the read-sequence cache where possible; reads for which the full sequence cannot be retrieved are excluded from the trimmed read set. The read interval retained for assembly is the maximal interval required by all connected candidate regions that the read supports. This prevents a read that spans multiple nearby signals from being cut separately at each signal. For clipped reads, the retained buffer is the larger of the configured clipped-sequence buffer and the maximum insertion size in the candidate-region container. Each trimmed read stores its read and reference cut coordinates in its FASTA description for later padding and provenance.

Svirlpool then tries a fast clustering shortcut before running all-vs-all read alignment. For each read, it computes a two-dimensional feature vector $x_i = (I_i, D_i)$, where $I_i$ and $D_i$ are the summed insertion and deletion signal burdens observed for that read in the original candidate-region alignments. Reads with outlying sequence lengths are removed before this shortcut: read lengths are sorted, a new length group is started whenever adjacent lengths differ by a factor greater than 1.2, and length groups with fewer than $\max(2, \lfloor\sqrt{n/\max(k,1)}\rfloor)$ reads are treated as outliers. KMeans is then evaluated for $k = 1, \ldots, \mathrm{CN}$, where CN is the local copy-number bound. Clustering uses `n_init=10` and `random_state=42`. For $k = 1$, the cluster is accepted only if the mean Euclidean distance of reads to the centroid is at most 14.5 bp. For $k > 1$, the maximum mean within-cluster distance must be at most 29 bp and the minimum Euclidean distance between any two cluster centroids must be at least 29 bp. If no $k$ satisfies these criteria, Svirlpool falls back to the all-vs-all alignment pathway.

In the fallback pathway, Svirlpool writes all trimmed reads in the container to a FASTA file and aligns the reads against each other with minimap2 in AVA ONT mode, using `--sam-hit-only --secondary=yes -U 25,35 -H`. Up to four AVA attempts are made. If an AVA attempt fails or produces no alignments, the read pool is deterministically subsampled to half its current size using seeds derived from 42, with at least two reads retained; excluded reads are later treated as isolated. Insertion and deletion signals of at least 12 bp and break-end signals of at least 100 bp are parsed from the directed AVA alignments.

The AVA fallback constructs a read-read similarity matrix from directed alignment signals. Small indel signals are damped by the monotonic function

$$g(s) = 1 - \exp\left[-\left(\frac{s}{30}\right)^{1.5}\right],$$

where $s$ is the absolute signal size in base pairs. This assigns little weight to very small indels and approaches one for larger events. For insertion and deletion signals, Svirlpool can also build size-density tracks: each observed signal size adds a smooth sigmoid plateau centered on the signal size, with a half-window of 20% of that size and sigmoid falloff 0.5. Positional importance-density tracks are built per read by adding a smooth 100 bp plateau centered on each non-BND signal, also with falloff 0.5. Size-density scaling is normalized per read and raised to power 1.5 before being applied to the positional density. The CLI parameter `--densities-weight` controls whether these density values affect the pairwise weights; a value of zero gives flat density weights.

For an alignment in which read $A$ is the query and read $B$ is the reference, the directed signal distance is

$$d(A \to B) = \sum_{s \in S_{A\to B}} w_B(s)\, w_A(s)\, g(|\Delta_s|)\, |\Delta_s|,$$

where $S_{A\to B}$ contains the non-BND insertion and deletion signals, $\Delta_s$ is the signal size, and $w_A(s)$ and $w_B(s)$ are the density-derived weights at the signal positions on the query and reference reads. For a read pair, the symmetric signal distance uses the larger of the two available directed distances so that a signal-free reverse alignment cannot mask a signal-rich forward alignment. A length penalty is then added:

$$D(A,B) = \max\{d(A \to B), d(B \to A)\} + \lambda\left(1 - \frac{\min(L_A, L_B)}{\max(L_A, L_B)}\right),$$

where $L_A$ and $L_B$ are the read lengths. Unless specified otherwise, $\lambda$ is calibrated to the median finite non-zero signal distance. The same median is used as the Gaussian kernel width $\sigma$, and the final similarity is

$$\mathrm{sim}(A, B) = \exp\left(-\frac{D(A, B)^2}{2\sigma^2}\right).$$

Read pairs with no AVA alignment have similarity zero, the diagonal is set to one, and any pair with a BND more than 100 bp from either end of a read has its similarity set to zero.

Before spectral clustering, Svirlpool removes outlier reads from the similarity matrix. Length outliers are identified with the same adjacent-length ratio rule used in the KMeans shortcut. Connectivity outliers are reads whose mean similarity to other reads is below the implementation threshold of 0.1. The remaining well-connected reads are clustered by scikit-learn `SpectralClustering` with a precomputed affinity matrix, `assign_labels="kmeans"`, and `random_state=42`. The requested number of clusters is the local copy-number bound, reduced when fewer well-connected reads remain. If all clustered consensus assemblies fail, isolated reads are assembled together as a rescue attempt.

For each KMeans or spectral-clustering read cluster, Svirlpool assembles a local consensus. In the benchmarked workflow described in the main text this was done with lamassemble, using `lamassemble --name <ID> -P <threads> -f fa -s 2 -g 67 <matrix> <reads>`. Assembly is run under the configured timeout. The code also contains alternative consensus backends based on racon or a Gotoh multiple-sequence alignment implementation, but these are not the lamassemble pathway described in the main manuscript.

After assembly, the trimmed reads are aligned back to the consensus with minimap2 using `-Y --sam-hit-only --secondary=no -U 10,25 -H`. Svirlpool records the alignment interval of each trimmed read on the consensus and parses insertion, deletion, and break-end distortions from these cut-read-to-consensus alignments, using minimum sizes of 8 bp for indels and 100 bp for break ends. These stored distortions are later used as the local read-derived noise information for variant scoring and merging.

Finally, Svirlpool stores both the consensus core and a padded consensus sequence. The core is the sequence produced from the trimmed reads. To make this local sequence easier to map, Svirlpool selects the supporting read with the largest left overhang and the supporting read with the largest right overhang, extracts the corresponding flanking sequence from the original full-length reads, and reverse complements the flank if the read aligned in the reverse orientation. The padded sequence is stored as lower-case left flank, upper-case consensus core, and lower-case right flank. The coordinates of the upper-case core within the padded sequence are stored explicitly, so downstream SV extraction can distinguish the assembled consensus sequence from the flanking padding.

### S2 Supplementary Figures

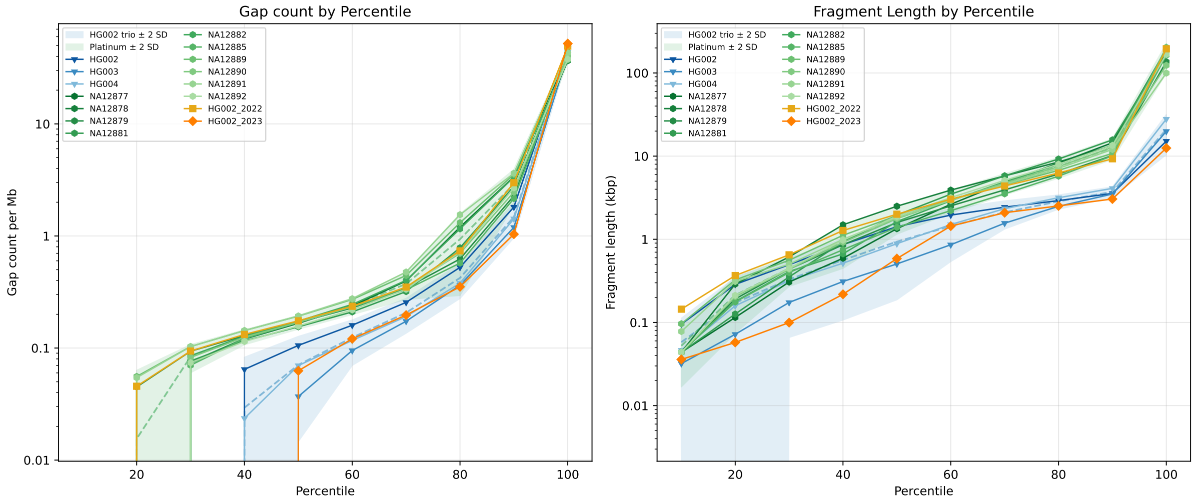


Figure S1: Gap counts and fragment lengths of the 30x HG002 trio ONT alignments, the platinum pedigree ONT alignments, and the 30x HG002 2022 and 2023 variants to hg38. The colors and markers separate the families and variants. The solid color bands indicate the two standard deviations range of the platinum pedigree family and the HG002 2025 trio.

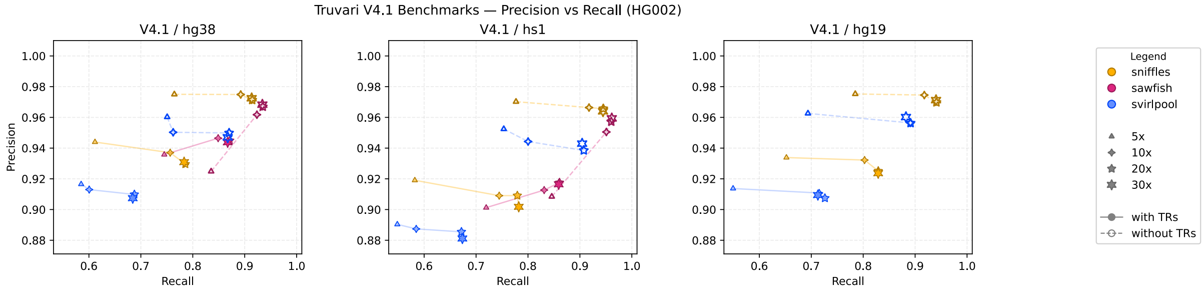


Figure S2: V4.1 benchmarks. We show the data separated by tandem repeat overlap (with TR vs. without TR). There are no results for HG19 as we faced technical challenges having Sawfish work with the HG19 reference that did not seem worthy to overcome, since the HG19 reference is considered outdated and not of much interest anymore.

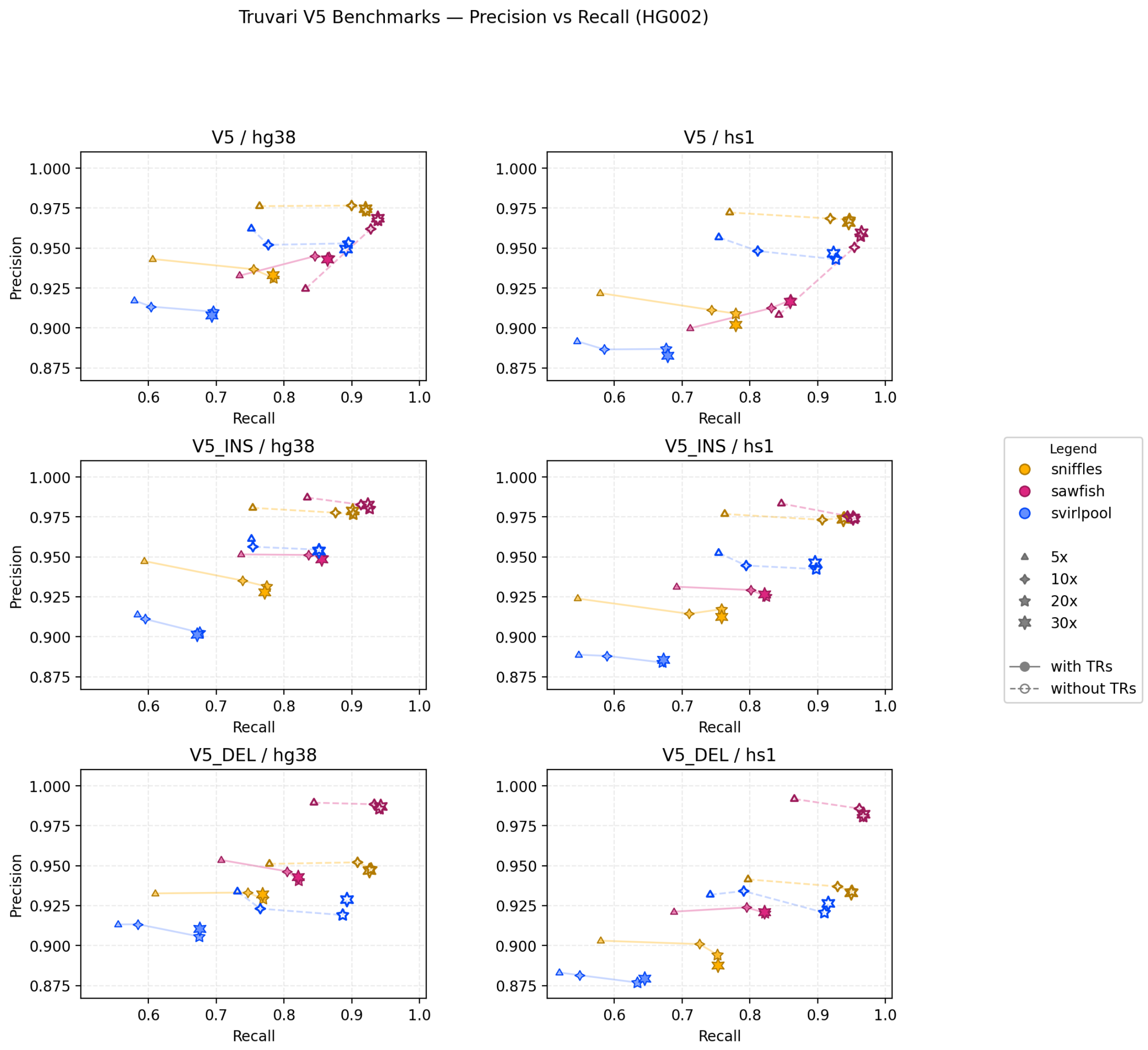


Figure S3: Complete v5 results, both with and without tandem repeats.

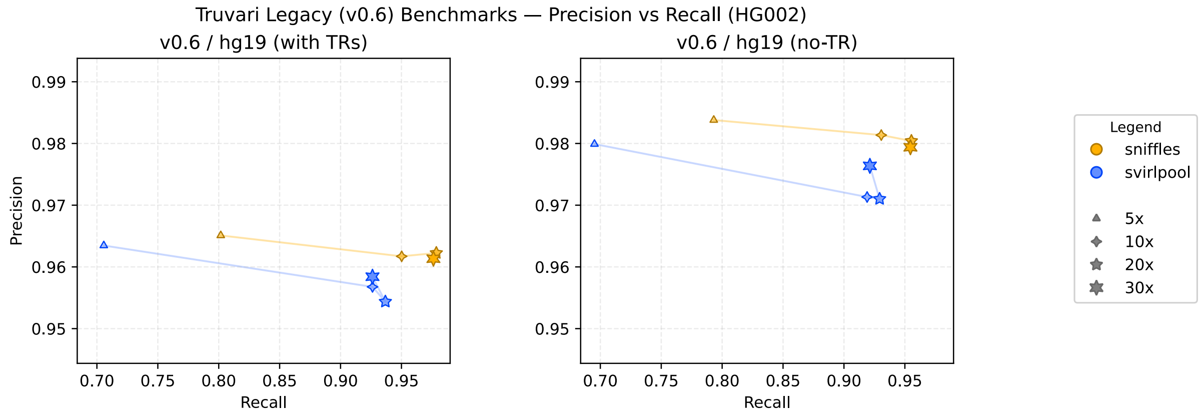


Figure S4: SV benchmark results on HG19. No split of with-TR and without-TR is necessary, as the benchmark respects the convention of left-annotation and merging SVs of the same allele within the same tandem repeat.

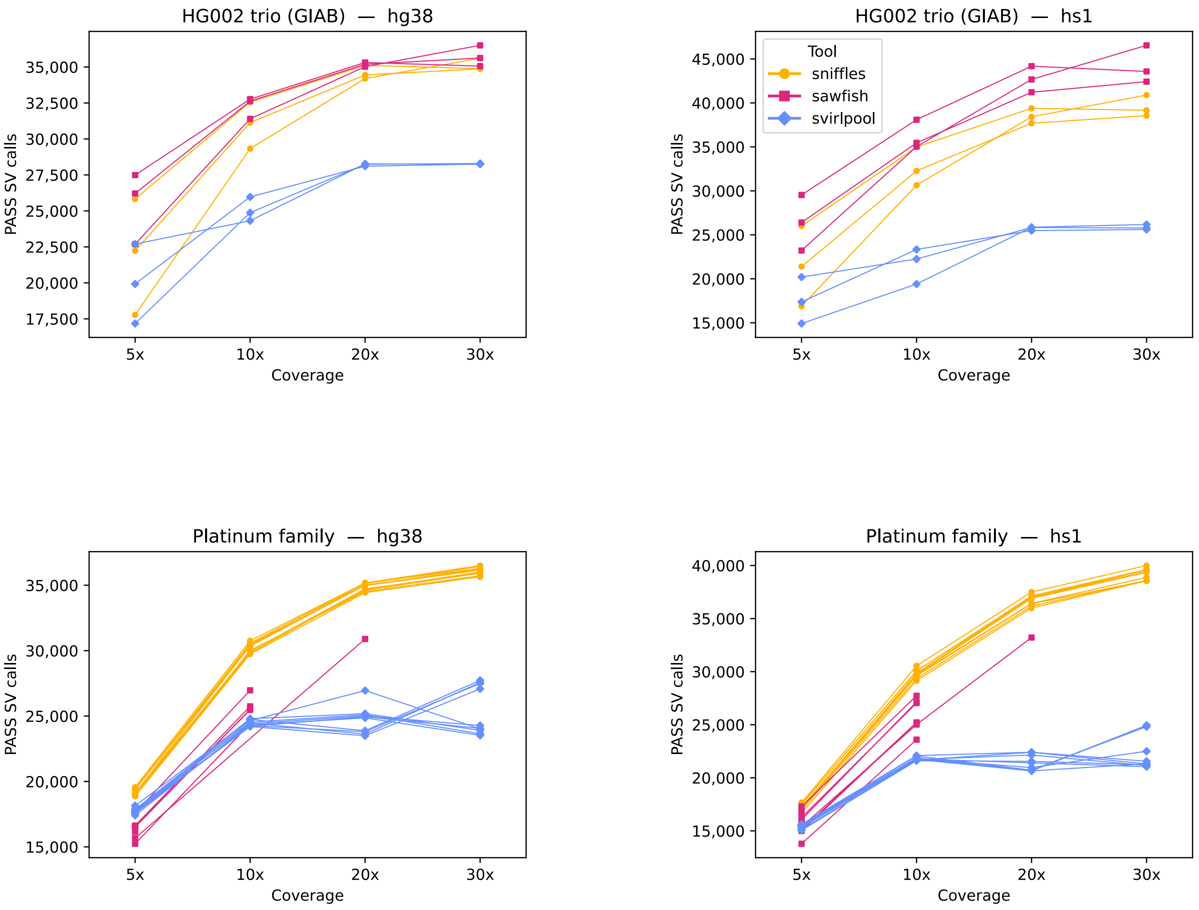


Figure S5: Absolute counts of called SVs that pass the quality filter (`FILTER=PASS`) by tool (color) and reference genome (sub-plot title), generated on the 30x single-sample data of HG002 trio and Platinum Pedigree samples. Variants smaller than 50 bp were excluded. Each group of bars corresponds to a coverage level (5x–30x); each bar shows one sample.

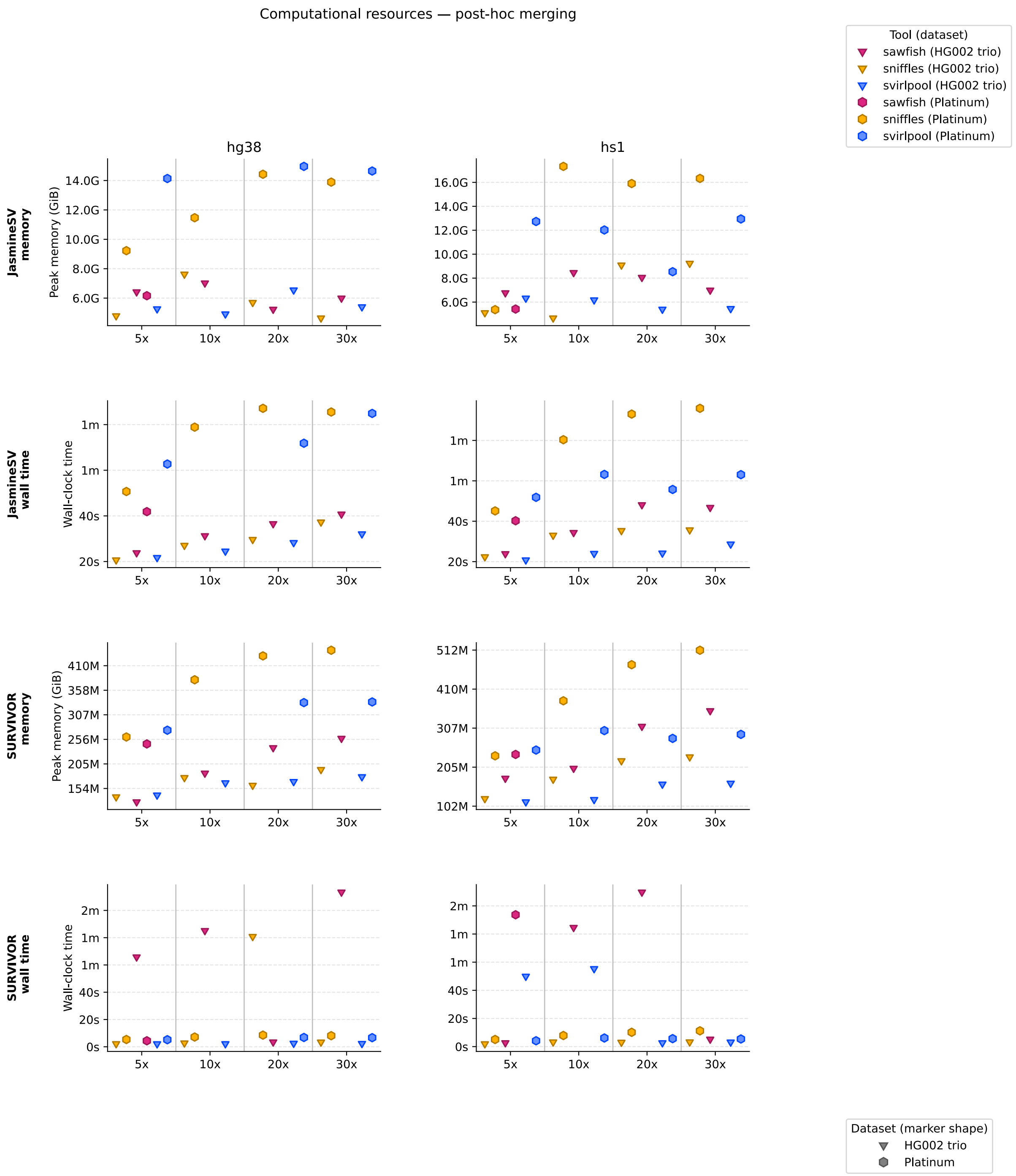


Figure S6: Post-hoc merging resource consumptions. Peak resident memory (GiB) and wall-clock time are shown for Jasmine and Survivor applied to the single-sample calls of each tool, stratified by reference genome (hg38 and hs1) and dataset (HG002 trio and Platinum Pedigree).

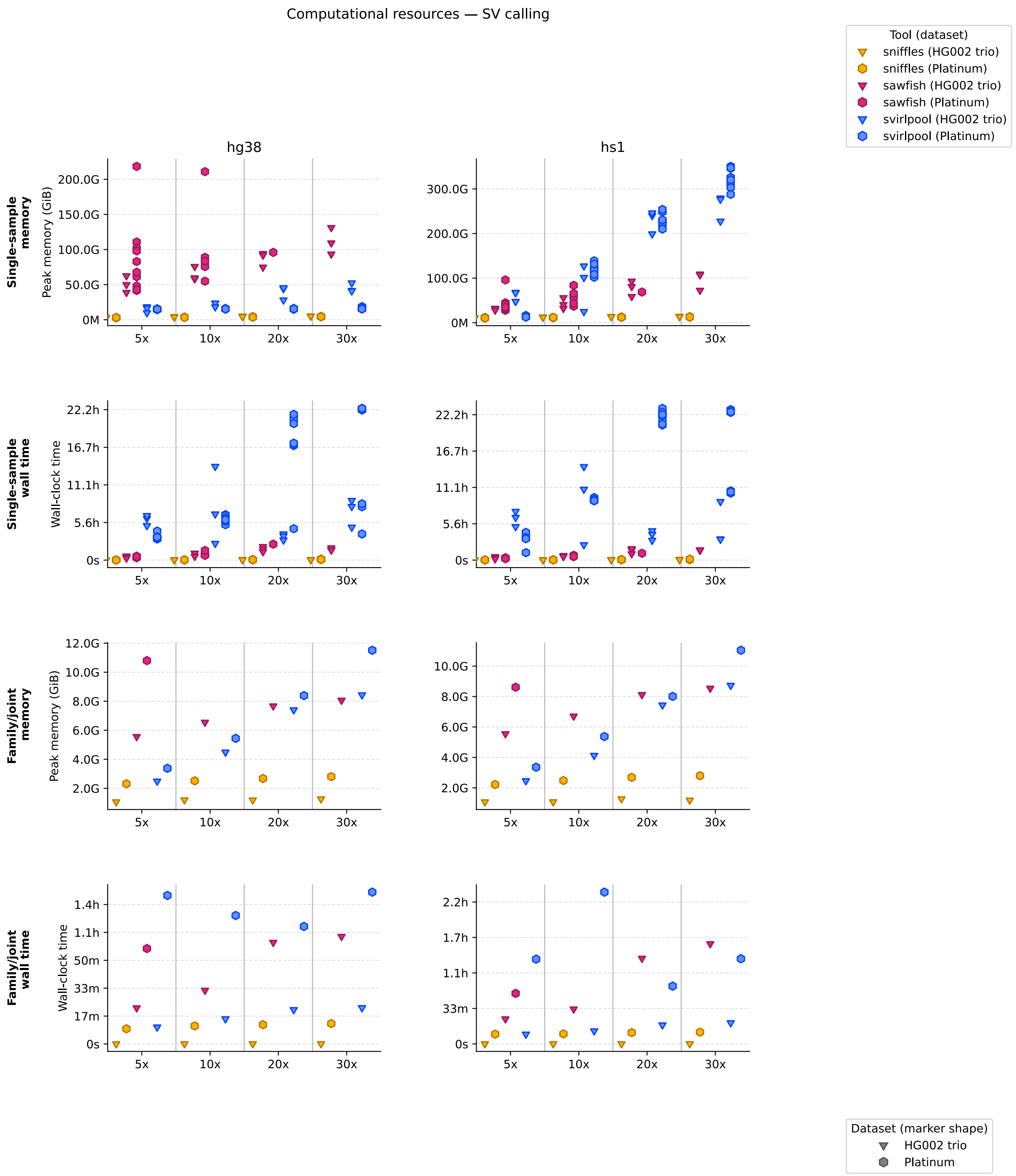


Figure S7: SV-caller resource consumptions. Peak resident memory (GiB) and wall-clock time for single-sample calling and family/joint calling are shown for Sniffles, Sawfish, and Svirlpool, stratified by reference genome (hg38 and hs1) and dataset (HG002 trio and Platinum Pedigree).

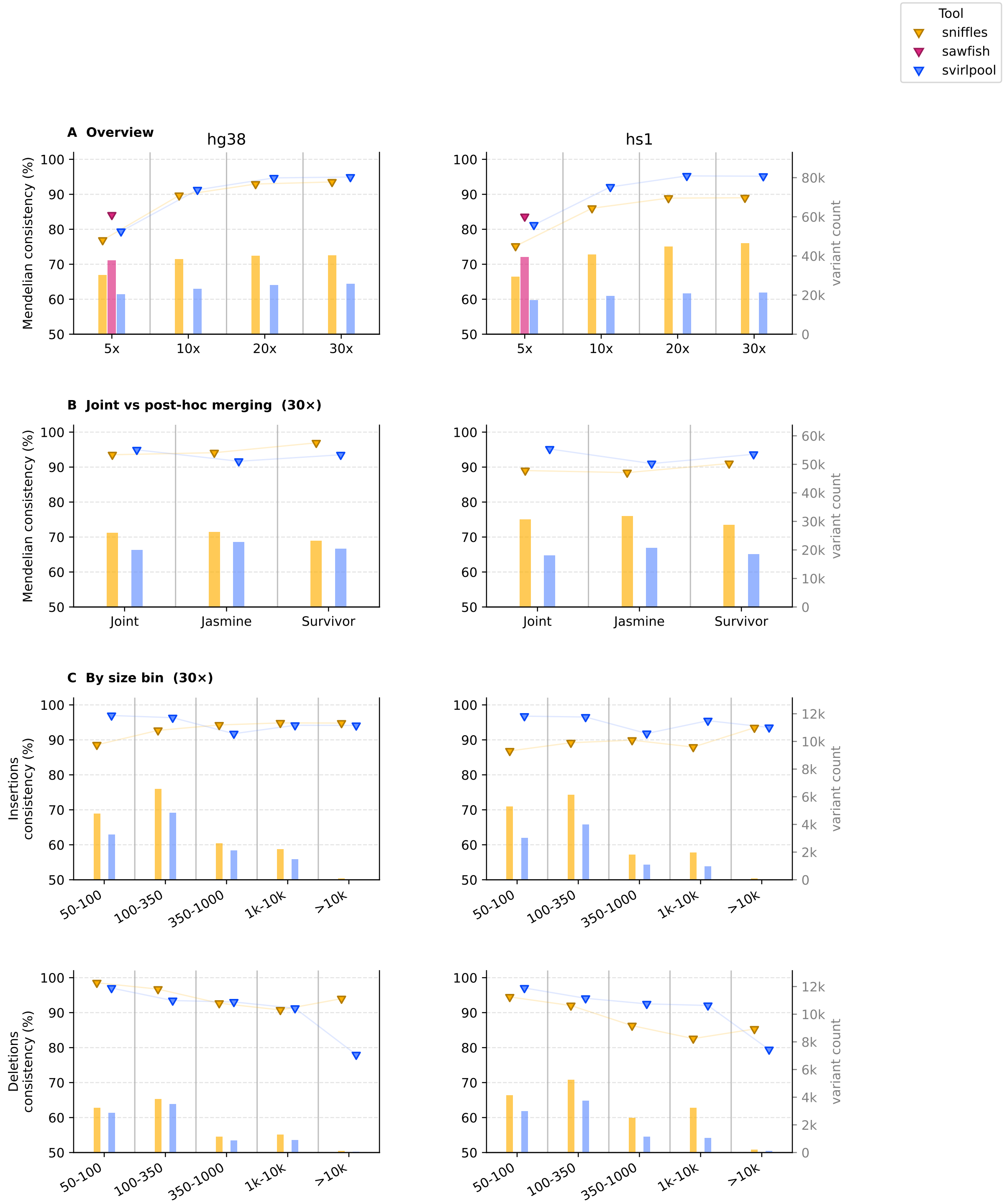


Figure S8: Mendelian consistency of a mixed-chemistry HG002 trio (child from the 2022 R9.4.1 or 2023 R10.4.1 dataset; parents from the 2025 HG002 trio). The layout mirrors Fig. 2 of the main text. **Sub-figure A**: consistency stratified by coverage (5x–30x) on GRCh38 (left) and CHM13v2 (right), with variant counts shown as bars. **Sub-figure B**: comparison of native joint calling against post-hoc Jasmine and Survivor merging at 30x. **Sub-figure C**: 30x results stratified by SV type (insertions, top; deletions, bottom) and size bin.

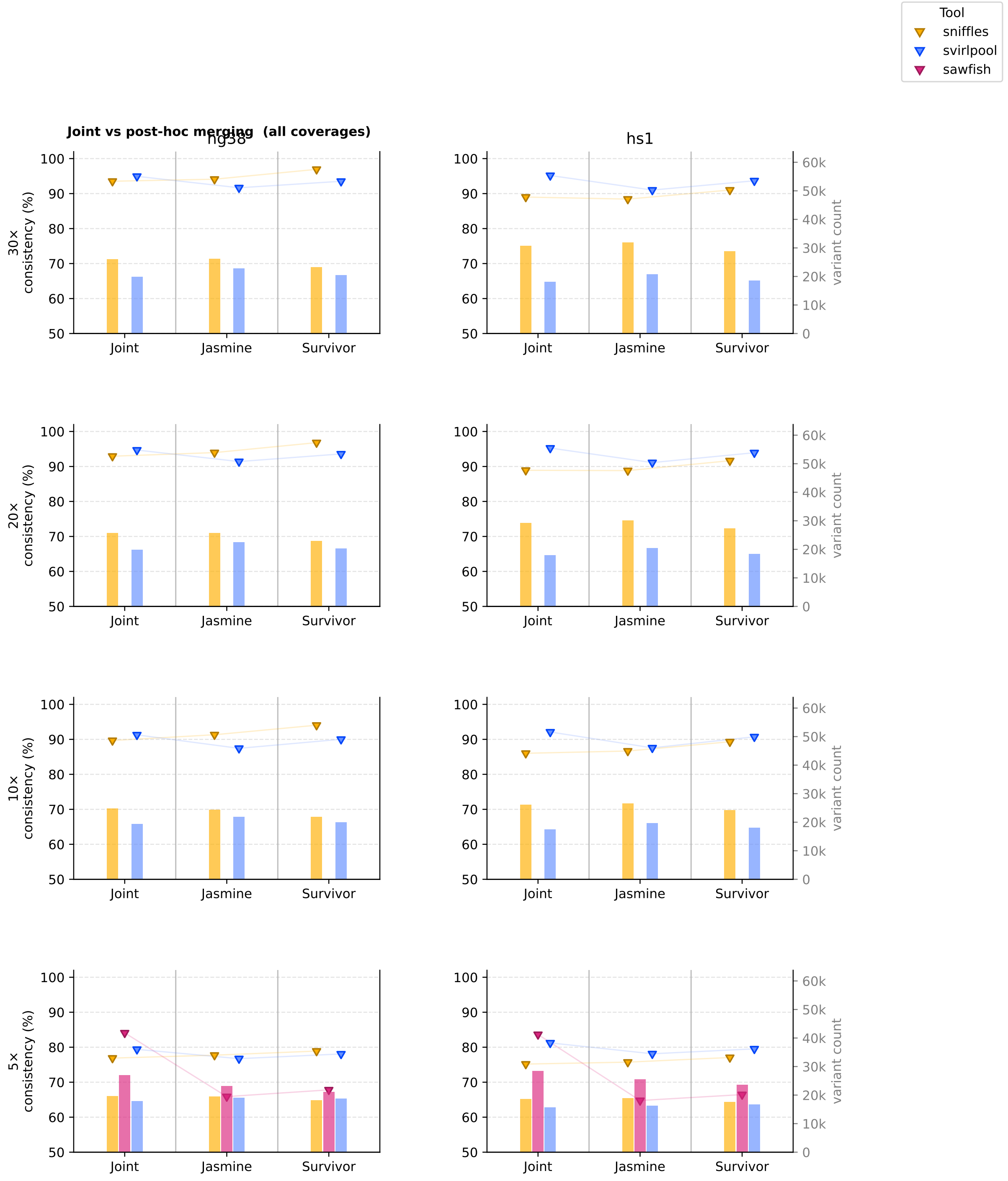


Figure S9: Joint calling vs. post-hoc merging (Jasmine, Survivor) for the mixed-chemistry HG002 batch experiment across all coverage levels (5x, 10x, 20x, 30x; rows) and both reference genomes (GRCh38, left; CHM13v2, right). Mendelian consistency (markers, left axis) and variant counts (bars, right axis) are shown for each combination of tool and merging strategy.

#### S3 Supplementary Figures S10–S13: IGV examples

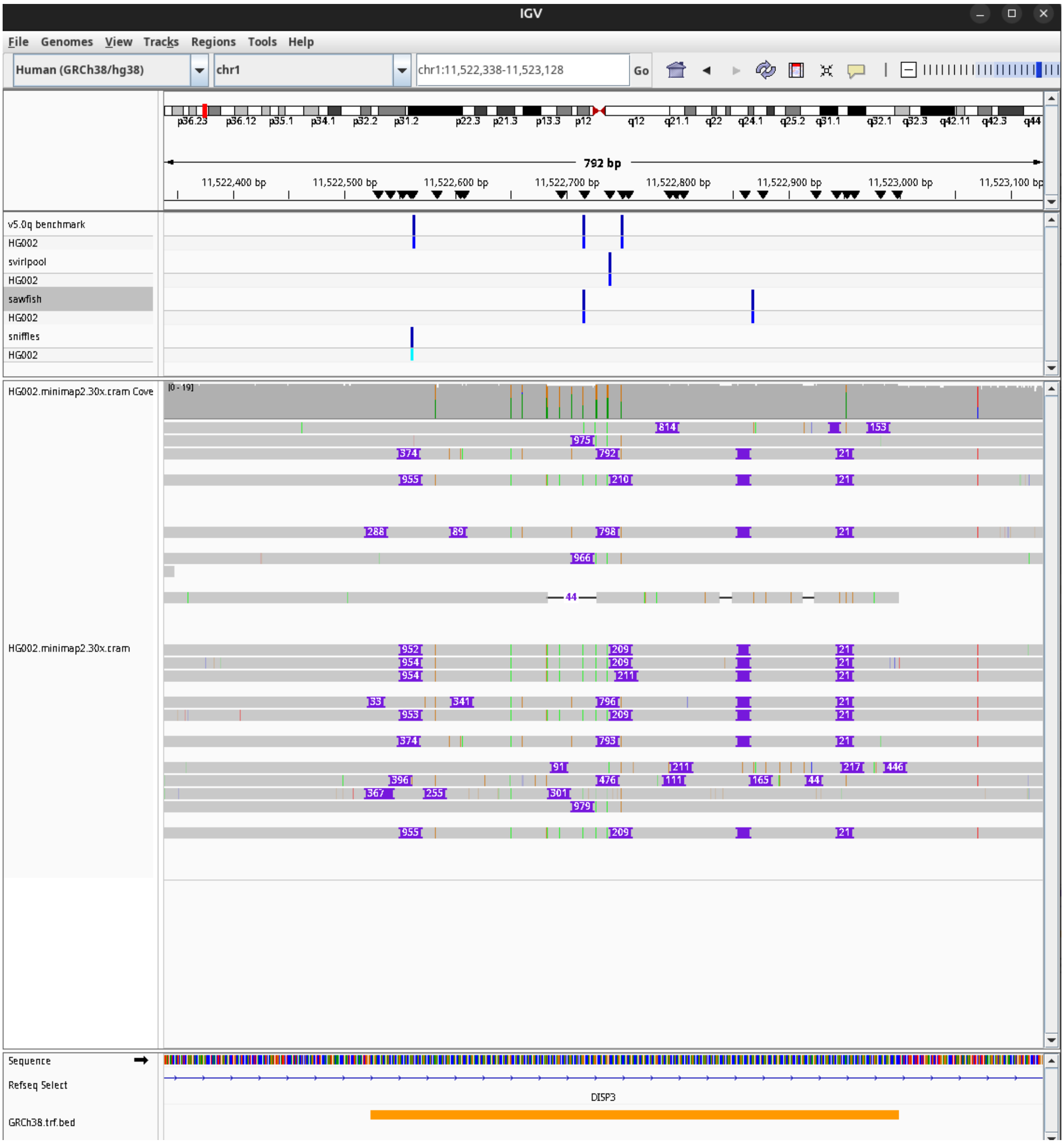


Figure S10: IGV screenshot. The orange track at the bottom shows tandem repeat annotations. The repeat expansion causes three SV calls in the SV benchmark, while Sawfish reports two insertions, Sniffles and Svirlpool one insertion.

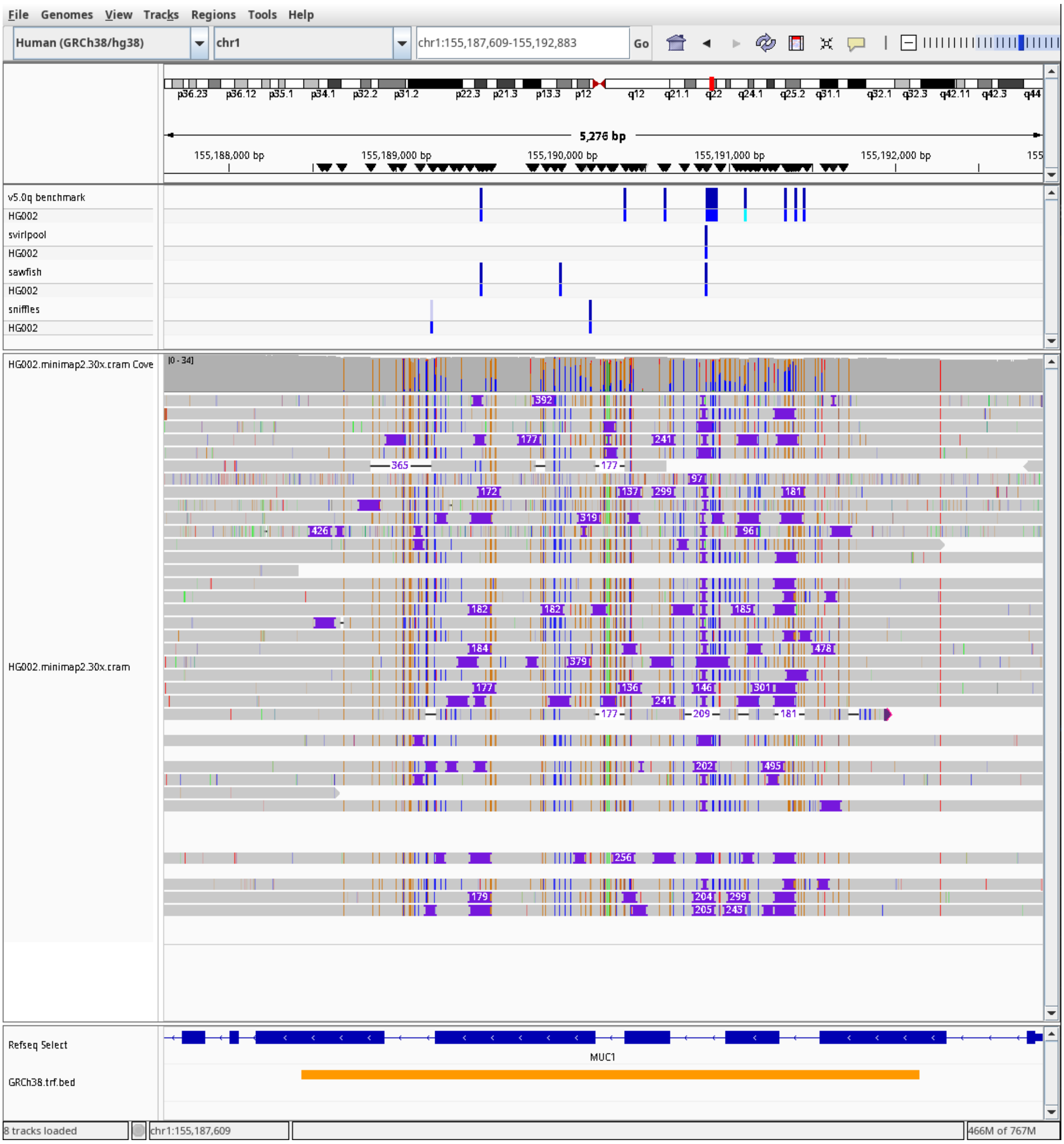


Figure S11: IGV screenshot. The orange track at the bottom shows tandem repeat annotations. The benchmark features seven insertions and one deletion, all larger than 50bp. Sawfish boils them down to three insertions, and both Sniffles and Svirlpool both report one. Note the genotypes: (blue is het and teal is hom) All three tools only identify heterozygous variants.

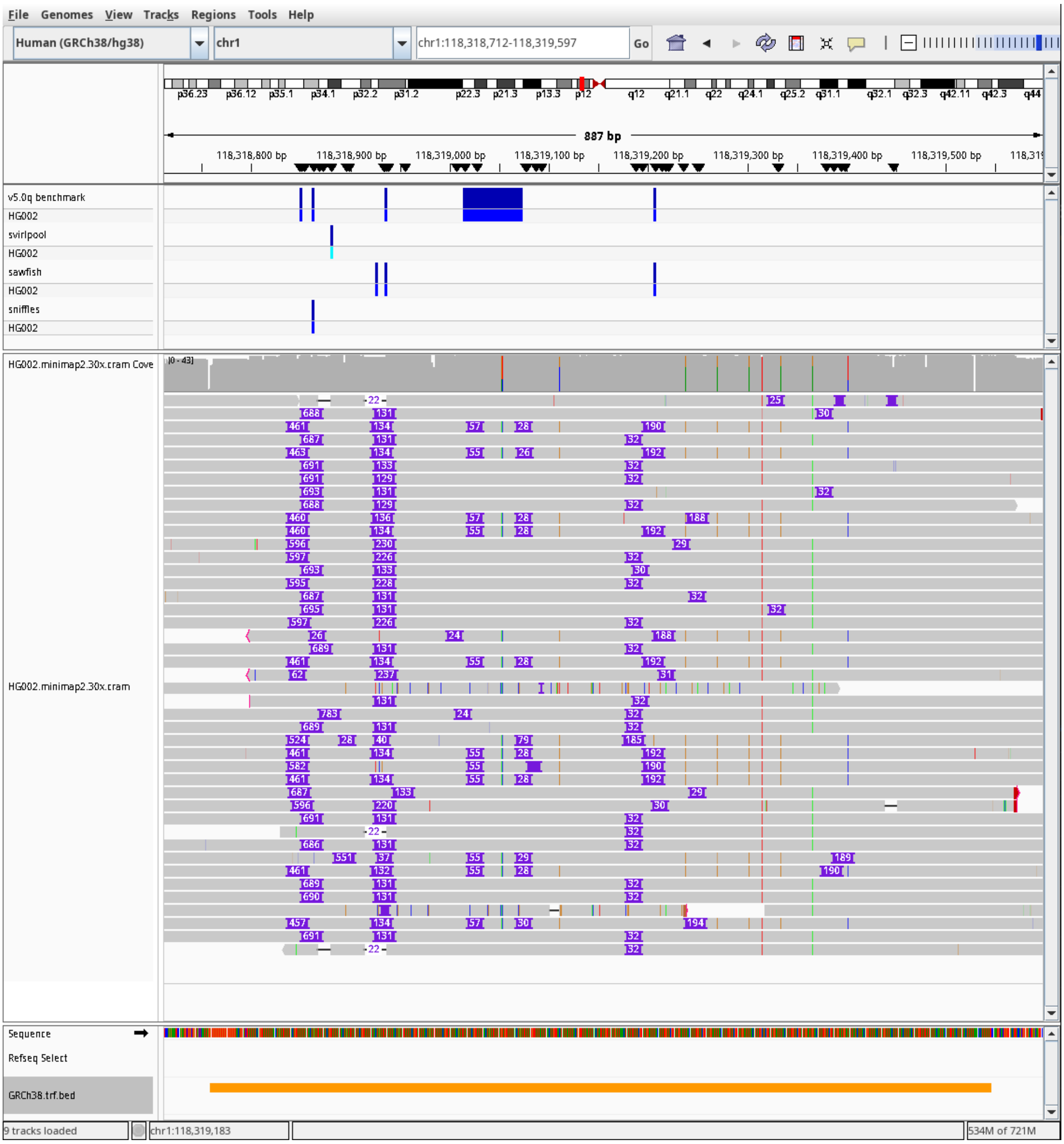


Figure S12: IGV screenshot. The orange track at the bottom shows tandem repeat annotations. The benchmark features four insertions and one deletion of 63bp. No other tool reports a deletion. Sawfish reports three insertions, and Sniffles and Svirlpool both agree about finding one. Note the genotype: Svirlpool identifies a homozygous variant.

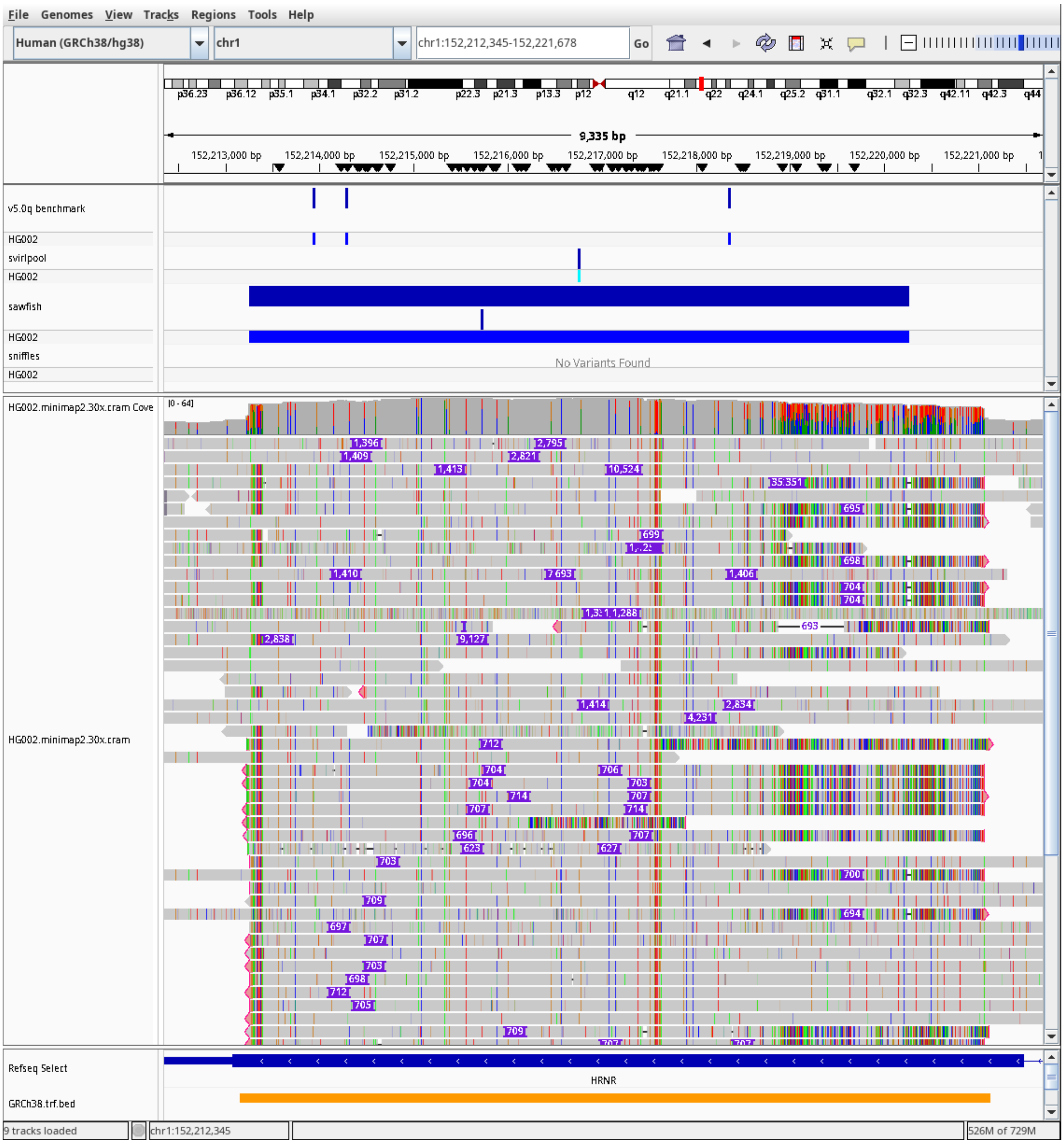


Figure S13: IGV screenshot. The benchmark reports three insertions here, while Sawfish identifies one het duplication (large blue bar), and one het insertion (small blue stripe). Svirlpool merges the haplotypes into one homozygous insertion. Sniffles misses the call.

#### S4 Supplementary Tables

Table S1: Per-run single-sample resource consumption (wall time and peak resident memory) for all tools, coverage levels, reference genomes, and datasets.

| Tool | Dataset | Reference | Coverage | Wall time (min) | | | Peak memory (GB) | | |
|---|---|---|---|---|---|---|---|---|---|
| | | | | Min | Median | Max | Min | Median | Max |
| Sawfish | Platinum | hg38 | 5x | 21.4 | 28.2 | 34.7 | 41.7 | 75.3 | 218.4 |
| Sawfish | Platinum | hg38 | 10x | 42.3 | 45.3 | 85.5 | 55.0 | 83.4 | 210.9 |
| Sawfish | Platinum | hg38 | 20x | 141.8 | 141.8 | 141.8 | 95.9 | 95.9 | 95.9 |
| Sniffles | Platinum | hg38 | 5x | 1.1 | 1.2 | 1.8 | 2.7 | 3.1 | 3.3 |
| Sniffles | Platinum | hg38 | 10x | 1.8 | 2.0 | 2.2 | 3.3 | 3.6 | 3.7 |
| Sniffles | Platinum | hg38 | 20x | 3.5 | 4.0 | 5.8 | 3.8 | 3.9 | 4.4 |
| Sniffles | Platinum | hg38 | 30x | 5.3 | 5.8 | 9.7 | 4.2 | 4.4 | 4.8 |
| Svirlpool | Platinum | hg38 | 5x | 187.6 | 201.6 | 257.9 | 14.4 | 14.9 | 15.6 |
| Svirlpool | Platinum | hg38 | 10x | 314.3 | 370.9 | 402.9 | 15.2 | 15.7 | 15.9 |
| Svirlpool | Platinum | hg38 | 20x | 278.5 | 1214.2 | 1291.9 | 15.0 | 15.6 | 16.1 |
| Svirlpool | Platinum | hg38 | 30x | 231.6 | 1331.0 | 1345.4 | 15.4 | 15.7 | 18.6 |
| Sawfish | Platinum | hs1 | 5x | 13.0 | 18.7 | 21.3 | 27.8 | 37.9 | 95.9 |
| Sawfish | Platinum | hs1 | 10x | 32.3 | 33.5 | 43.9 | 36.9 | 56.6 | 83.7 |
| Sawfish | Platinum | hs1 | 20x | 62.7 | 62.7 | 62.7 | 68.7 | 68.7 | 68.7 |
| Sniffles | Platinum | hs1 | 5x | 0.9 | 1.1 | 1.3 | 10.4 | 10.6 | 12.0 |
| Sniffles | Platinum | hs1 | 10x | 1.8 | 2.3 | 3.3 | 11.2 | 11.7 | 12.0 |
| Sniffles | Platinum | hs1 | 20x | 3.6 | 3.9 | 5.4 | 12.0 | 12.3 | 12.7 |
| Sniffles | Platinum | hs1 | 30x | 4.4 | 6.3 | 9.2 | 12.6 | 12.9 | 13.3 |
| Svirlpool | Platinum | hs1 | 5x | 67.9 | 202.2 | 254.2 | 12.7 | 13.9 | 16.1 |
| Svirlpool | Platinum | hs1 | 10x | 543.5 | 554.7 | 573.4 | 102.0 | 118.1 | 138.7 |
| Svirlpool | Platinum | hs1 | 20x | 1239.0 | 1329.0 | 1392.2 | 210.1 | 227.7 | 253.8 |
| Svirlpool | Platinum | hs1 | 30x | 614.8 | 1358.2 | 1379.9 | 287.8 | 322.9 | 350.5 |
| Sawfish | Trio | hg38 | 5x | 17.4 | 29.5 | 35.4 | 38.8 | 49.9 | 62.6 |
| Sawfish | Trio | hg38 | 10x | 33.5 | 59.9 | 61.3 | 58.1 | 59.6 | 75.8 |
| Sawfish | Trio | hg38 | 20x | 73.8 | 96.4 | 119.3 | 74.8 | 91.9 | 94.1 |
| Sawfish | Trio | hg38 | 30x | 87.5 | 91.1 | 108.2 | 93.5 | 109.3 | 131.2 |
| Sniffles | Trio | hg38 | 5x | 1.2 | 1.7 | 1.8 | 2.4 | 3.0 | 3.6 |
| Sniffles | Trio | hg38 | 10x | 2.2 | 2.3 | 2.6 | 3.8 | 4.2 | 4.3 |
| Sniffles | Trio | hg38 | 20x | 2.8 | 3.3 | 5.2 | 4.4 | 4.7 | 5.1 |
| Sniffles | Trio | hg38 | 30x | 3.9 | 3.9 | 3.9 | 5.1 | 5.1 | 5.3 |
| Svirlpool | Trio | hg38 | 5x | 305.0 | 369.6 | 394.1 | 9.9 | 16.4 | 18.4 |
| Svirlpool | Trio | hg38 | 10x | 148.1 | 408.2 | 830.0 | 18.3 | 23.5 | 23.7 |
| Svirlpool | Trio | hg38 | 20x | 177.7 | 212.9 | 231.2 | 28.1 | 44.3 | 45.4 |
| Svirlpool | Trio | hg38 | 30x | 291.4 | 474.8 | 528.0 | 40.9 | 41.7 | 52.4 |
| Sawfish | Trio | hs1 | 5x | 10.8 | 19.4 | 30.7 | 28.0 | 31.3 | 32.1 |
| Sawfish | Trio | hs1 | 10x | 31.2 | 38.8 | 41.2 | 31.9 | 40.4 | 56.0 |
| Sawfish | Trio | hs1 | 20x | 56.9 | 92.1 | 103.9 | 58.7 | 81.0 | 92.9 |
| Sawfish | Trio | hs1 | 30x | 89.3 | 91.1 | 93.8 | 72.5 | 107.1 | 108.8 |
| Sniffles | Trio | hs1 | 5x | 0.9 | 1.0 | 1.1 | 10.5 | 10.9 | 11.0 |
| Sniffles | Trio | hs1 | 10x | 1.6 | 1.8 | 2.1 | 12.1 | 12.3 | 12.6 |
| Sniffles | Trio | hs1 | 20x | 3.4 | 3.7 | 4.3 | 13.2 | 13.6 | 13.8 |
| Sniffles | Trio | hs1 | 30x | 4.5 | 5.0 | 5.1 | 13.4 | 13.7 | 14.1 |
| Svirlpool | Trio | hs1 | 5x | 306.3 | 391.0 | 446.3 | 47.4 | 67.1 | 67.9 |
| Svirlpool | Trio | hs1 | 10x | 140.3 | 649.0 | 856.5 | 24.9 | 101.3 | 127.1 |
| Svirlpool | Trio | hs1 | 20x | 181.0 | 235.8 | 268.5 | 199.0 | 240.2 | 246.0 |
| Svirlpool | Trio | hs1 | 30x | 190.1 | 193.9 | 536.5 | 227.6 | 276.5 | 279.4 |

Table S2: Resource consumption for the joint-calling step of all tools (Sniffles, Sawfish, Svirlpool, Jasmine, and Survivor).

| Tool | Dataset | Reference | Coverage | Wall time (min) | Peak memory (GB) |
|---|---|---|---|---|---|
| Sawfish | Platinum | hg38 | 5x | 57.0 | 10.8 |
| Sniffles | Platinum | hg38 | 5x | 9.1 | 2.3 |
| Svirlpool | Platinum | hg38 | 5x | 88.8 | 3.4 |
| Sniffles | Platinum | hg38 | 10x | 10.8 | 2.5 |
| Svirlpool | Platinum | hg38 | 10x | 76.8 | 5.4 |
| Sniffles | Platinum | hg38 | 20x | 11.5 | 2.7 |
| Svirlpool | Platinum | hg38 | 20x | 70.2 | 8.4 |
| Sniffles | Platinum | hg38 | 30x | 12.2 | 2.8 |
| Svirlpool | Platinum | hg38 | 30x | 90.7 | 11.5 |
| Sawfish | Platinum | hs1 | 5x | 47.5 | 8.6 |
| Sniffles | Platinum | hs1 | 5x | 9.2 | 2.2 |
| Svirlpool | Platinum | hs1 | 5x | 79.7 | 3.4 |
| Sniffles | Platinum | hs1 | 10x | 9.4 | 2.5 |
| Svirlpool | Platinum | hs1 | 10x | 142.6 | 5.4 |
| Sniffles | Platinum | hs1 | 20x | 10.6 | 2.7 |
| Svirlpool | Platinum | hs1 | 20x | 54.3 | 8.0 |
| Sniffles | Platinum | hs1 | 30x | 11.1 | 2.8 |
| Svirlpool | Platinum | hs1 | 30x | 80.0 | 11.0 |
| Sawfish | Trio | hg38 | 5x | 21.5 | 5.6 |
| Sniffles | Trio | hg38 | 5x | 0.1 | 1.1 |
| Svirlpool | Trio | hg38 | 5x | 10.1 | 2.5 |
| Sawfish | Trio | hg38 | 10x | 32.0 | 6.5 |
| Sniffles | Trio | hg38 | 10x | 0.1 | 1.2 |
| Svirlpool | Trio | hg38 | 10x | 15.0 | 4.5 |
| Sawfish | Trio | hg38 | 20x | 60.7 | 7.7 |
| Sniffles | Trio | hg38 | 20x | 0.1 | 1.2 |
| Svirlpool | Trio | hg38 | 20x | 20.5 | 7.4 |
| Sawfish | Trio | hg38 | 30x | 64.2 | 8.1 |
| Sniffles | Trio | hg38 | 30x | 0.1 | 1.3 |
| Svirlpool | Trio | hg38 | 30x | 21.6 | 8.4 |
| Sawfish | Trio | hs1 | 5x | 23.5 | 5.6 |
| Sniffles | Trio | hs1 | 5x | 0.1 | 1.1 |
| Svirlpool | Trio | hs1 | 5x | 9.1 | 2.5 |
| Sawfish | Trio | hs1 | 10x | 32.7 | 6.7 |
| Sniffles | Trio | hs1 | 10x | 0.1 | 1.1 |
| Svirlpool | Trio | hs1 | 10x | 12.2 | 4.1 |
| Sawfish | Trio | hs1 | 20x | 80.4 | 8.1 |
| Sniffles | Trio | hs1 | 20x | 0.1 | 1.3 |
| Svirlpool | Trio | hs1 | 20x | 17.8 | 7.4 |
| Sawfish | Trio | hs1 | 30x | 94.0 | 8.5 |
| Sniffles | Trio | hs1 | 30x | 0.1 | 1.2 |
| Svirlpool | Trio | hs1 | 30x | 19.8 | 8.7 |
